# The photosystem I supercomplex from a primordial green alga *Ostreococcus tauri* harbors three light-harvesting complex trimers

**DOI:** 10.1101/2022.11.08.515661

**Authors:** Asako Ishii, Jianyu Shan, Xin Sheng, Eunchul Kim, Akimasa Watanabe, Makio Yokono, Chiyo Noda, Chihong Song, Kazuyoshi Murata, Zhenfeng Liu, Jun Minagawa

## Abstract

As a ubiquitous picophytoplankton in the ocean and an early-branching green alga, *Ostreococcus tauri* is a model prasinophyte species for studying the functional evolution of the light-harvesting systems in photosynthesis. Here, we report the structure and function of the *O. tauri* photosystem I (PSI) supercomplex in the low light, where it expands its photon-absorbing capacity by assembling with the light-harvesting complexes I (LHCI) and a prasinophyte-specific light-harvesting complex (Lhcp). Its architecture exhibits hybrid features of the plant-type and the green algal-type PSI supercomplexes, consisting of a PSI core, a Lhca1-Lhca4-Lhca2-Lhca3 belt attached on one side and a Lhca5-Lhca6 heterodimer associated on the other side between PsaG and PsaH. Interestingly, nine Lhcp subunits, including one Lhcp1 monomer with a phosphorylated amino-terminal threonine and eight Lhcp2 monomers, oligomerize into three trimers and associate with PSI on the third side between Lhca6 and PsaK. The Lhcp1 phosphorylation and the light-harvesting capacity of PSI were subjected to reversible photoacclimation, suggesting that the formation of *Ot*PSI–LHCI–Lhcp supercomplex is likely due to a state transition-like mechanism induced by light intensity change. Notably, this supercomplex did not exhibit far-red peaks in the 77 K fluorescence spectra, which is possibly due to weak coupling of the chlorophyll *a*603-*a*609 pair in *Ot*Lhca1-4.

## INTRODUCTION

Phytoplankton are the major primary producers in the aquatic environments, and provide organic matter for marine food webs by converting solar energy into chemical energy through photosynthesis. As a member of natural phytoplankton living in the ocean, *Ostreococcus tauri* is a unicellular green alga widespread in marine environments and crucial for the aquatic ecosystem(Derelle et al., 2006; Palenik et al., 2007). It is also known as the smallest free-living eukaryote(Courties et al., 1994) and belongs to Prasinophyceae, a class of green algae at the near basal position in the evolution of green lineage and most closely related to the first green alga known as “ancestral green flagellate”(Lewis & McCourt, 2004).

A prasinophyte-specific Lhc protein named Lhcp, was found in *O. tauri*(Six et al., 2005; Swingley et al., 2010) and *Mantoniella squamata* (the other member of Prasinophyceae)(Jiao & Fawley, 1994; Schmitt et al., 1994). There are two classes of Lhcp (Lhcp1 and Lhcp2) in addition to the common LHC proteins, namely six Lhca proteins (Lhca1-6) and two minor monomeric Lhcb proteins (Lhcb4 and Lhcb5), serving as the peripheral antennae of *O. tauri* PSI and PSII(Six et al., 2005; Swingley et al., 2010). The Lhcp proteins form a highly abundant antenna complexes in *O. tauri*(Swingley et al., 2010), and the carotenoid composition of the *Ot*Lhcp complexes is largely different from those of LHCIs or light-harvesting complexes II (LHCIIs) in plants and other green algae (such as *Chlamydomonas reinhardtii* of the Chlorophyceae class)(Minagawa, 2009). While Lhcp, Lhcb (apoproteins of LHCII) and Lhca (apoproteins of LHCI) all belong to the LHC superfamily, Lhcp proteins form a separate clade with characteristics of an ancestral state of LHC proteins instead of belonging to the clades of Lhcb or Lhca(Six et al., 2005). Moreover, the organization of pigment molecules within the Lhcp complexes exhibits distinct features in comparison with plant LHCII according to a previous spectroscopic study(Goss et al., 2000).

Under low light (LL) conditions, the Lhcp complexes of *O. tauri* can assemble with the PSI–LHCI (A3 band) to form a larger PSI–LHCI–Lhcp supercomplex (A3L band), whereas the amount of A3L is greatly reduced under high-light conditions(Swingley et al., 2010). It remains unclear how the Lhcp complexes assemble with PSI and establish energy transfer pathways with the interfacial PSI subunits under LL conditions. The arrangement of various Chl and carotenoid molecules within the Lhcp complexes is also unknown and awaits to be analyzed through further studies.

## RESULTS AND DISCUSSION

### The PSI–LHCI–Lhcp supercomplex

To stabilize the photosynthetic supercomplexes from *O. tauri* during purification process, the detergent-solubilized thylakoid membrane was treated with amphipol A8-35 before sucrose density gradient (SDG) ultracentrifugation (Fig. 1a), according to the previous protocol successfully employed to stabilize the PSII supercomplex from *C. reinhardtii*(Watanabe et al., 2019). Pigment composition analysis of the A3L fraction demonstrates that the most abundant light-harvesting carotenoid is prasinoxanthin (Prx, which is unique to prasinophytes) in the sample, among other carotenoids of lower abundance, including dihydrolutein (Dlt) and micromonal (Figure 1—supplement 1c and Figure 1—table supplement 1). Although the typical Chls (such as Chl *a* and Chl *b*) and carotenoids (violaxanthin/Vio, 9’-cis-neoxanthin/Nex and β-carotene) found in green plants were also present in *O. tauri*, lutein (a major carotenoid found in plant LHCII) was not detected. Polypeptides for the PSI core and its peripheral antennae in the A3L fraction were characterized by SDS-PAGE (mass spectrometry analysis (Appendix 1-table 1). These results indicate that A3L (Figure 1—supplement 1b) is mainly composed of the PSI–LHCI–Lhcp supercomplex.

**Fig. 1.**
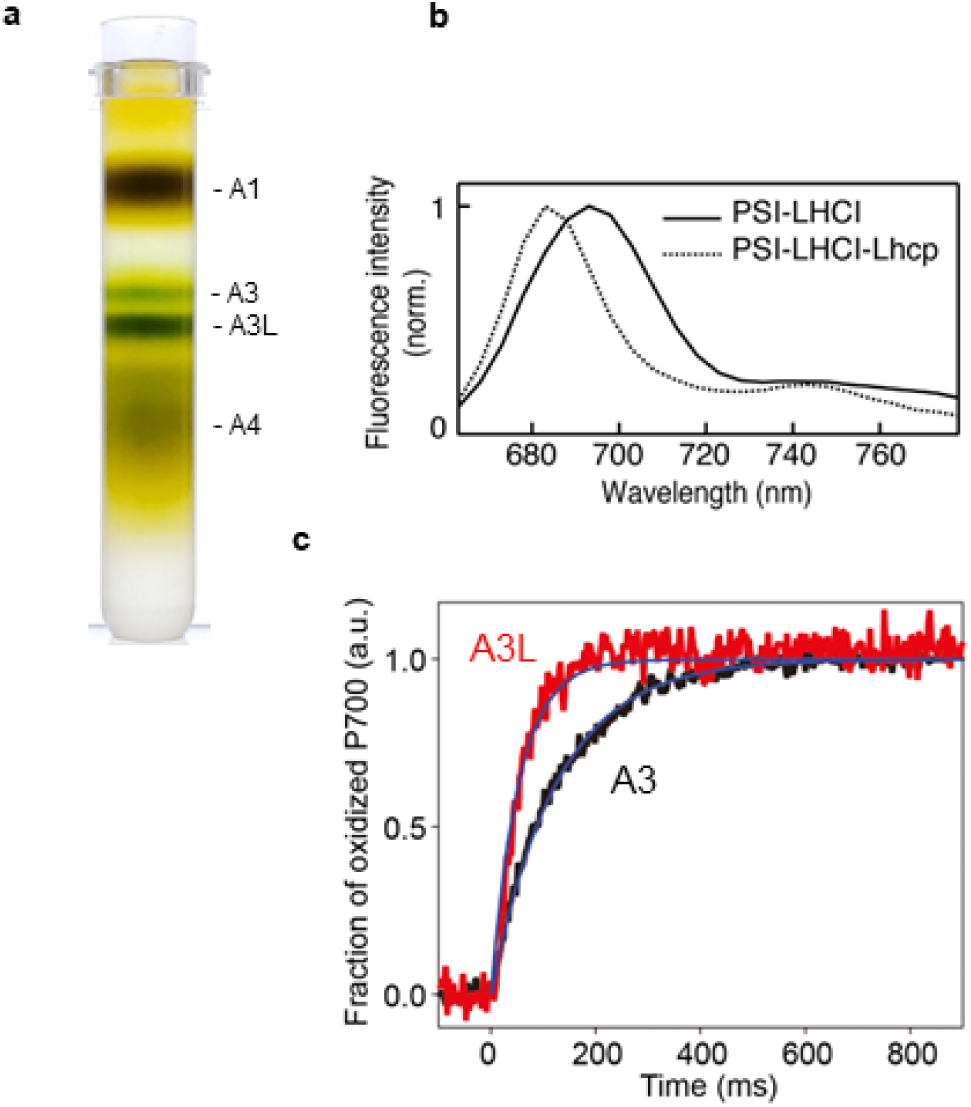
Characterization of A3 and A3L fractions. **a**, Sucrose density gradient showing four major bands corresponding to free LHCs (A1), PSI-LHCI supercomplex (A3), PSI-LHCI-Lhcp supercomplex, PSII-Lhcp supercomplex (A3L), and other complexes (A4). **b**, 77K steady-state fluorescence spectrum of PSI-LHCI (solid line) and PSI-LHCI-Lhcp (dotted line). **c**, PSI light-harvesting capabilities in the A3 and A3L fractions. Light-induced P700 oxidation kinetics of PSI were measured in A3 and A3L fractions under 28 µmol photon m^-2^s^-1^. The fraction of P700 oxidation was derived from Δ(A_820_-A_870_). *Solid* lines and *shaded* area represent averages of five (for A3) or ten (for A3L) technical replicates and SD, respectively. *Blue* lines represent fitting curves by mono-exponential functions. Data are representative of two biologically independent experiments. See another set of data in Figure 1—supplement 3. **Source data 1.** Quantitative data for Figure 1c. **Figure supplement 1.** Fractionation and characterization of the supercomplex samples from the *O. tauri* cells grown in the low light (50 µmol photon m^-2^ s^-1^). **Figure supplement 1—source data 1.** Raw data for Figure 1***—figure supplement 1***. **Figure supplement 2.** Fluorescence decay-associated spectra (FDAS) at 77K (ex., 405 nm, 4 µg Chl ml^-1^). **Figure supplement 3.** PSI light-harvesting capabilities in the A3 and A3L fractions. **Figure supplement 3—source data 1.** Quantitative data for Figure 1***—figure supplement 3***. **Figure supplement 4.** Structural analysis of A3 and A3L fractions by negative staining EM. **Table supplement 1.** Pigment composition in the A3L fraction as revealed by UPLC analysis. **Table supplement 2.** Two-dimensional classification of the PSI particles in the A3 and A3L fractions.

**Table 1.**
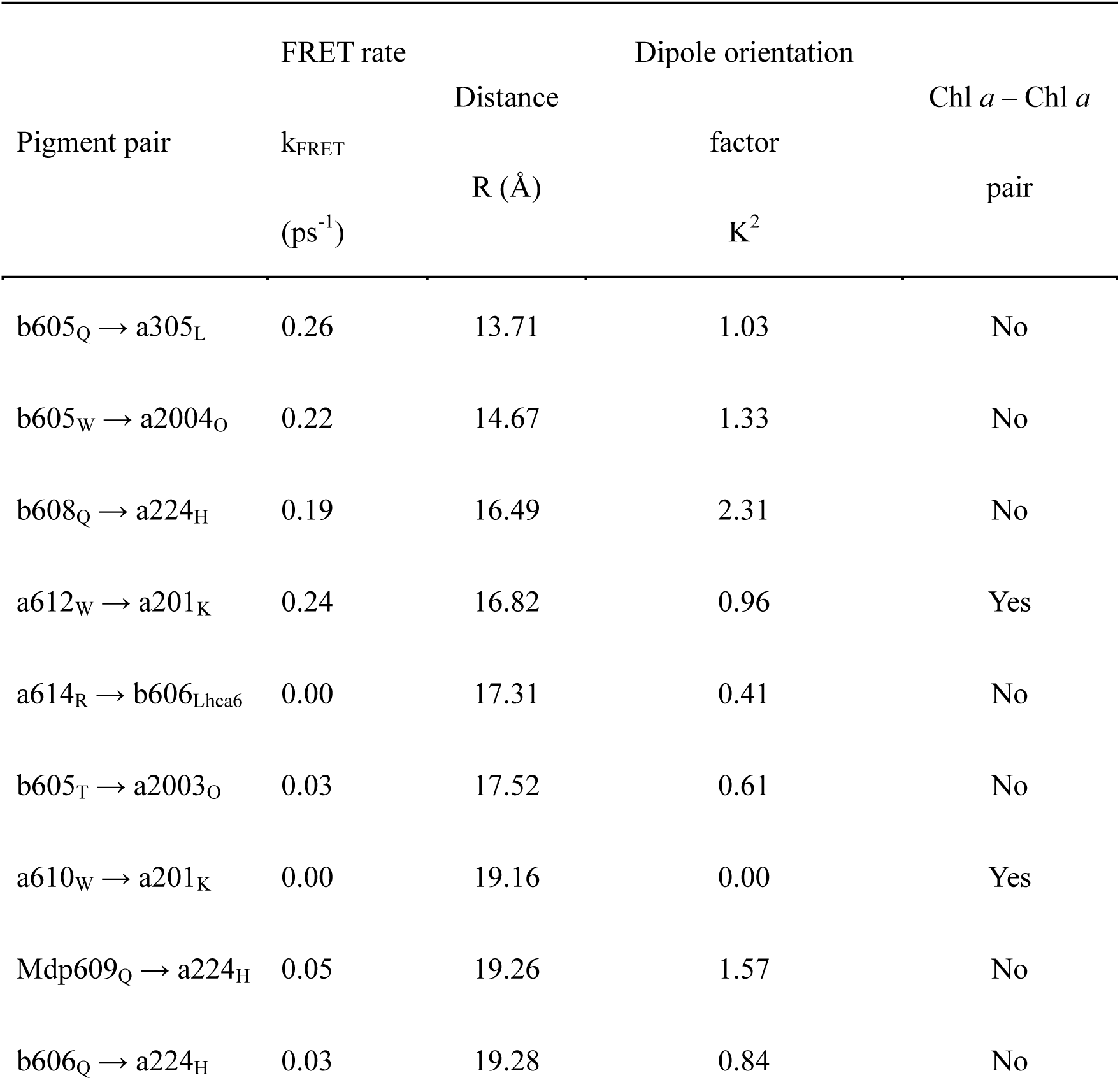
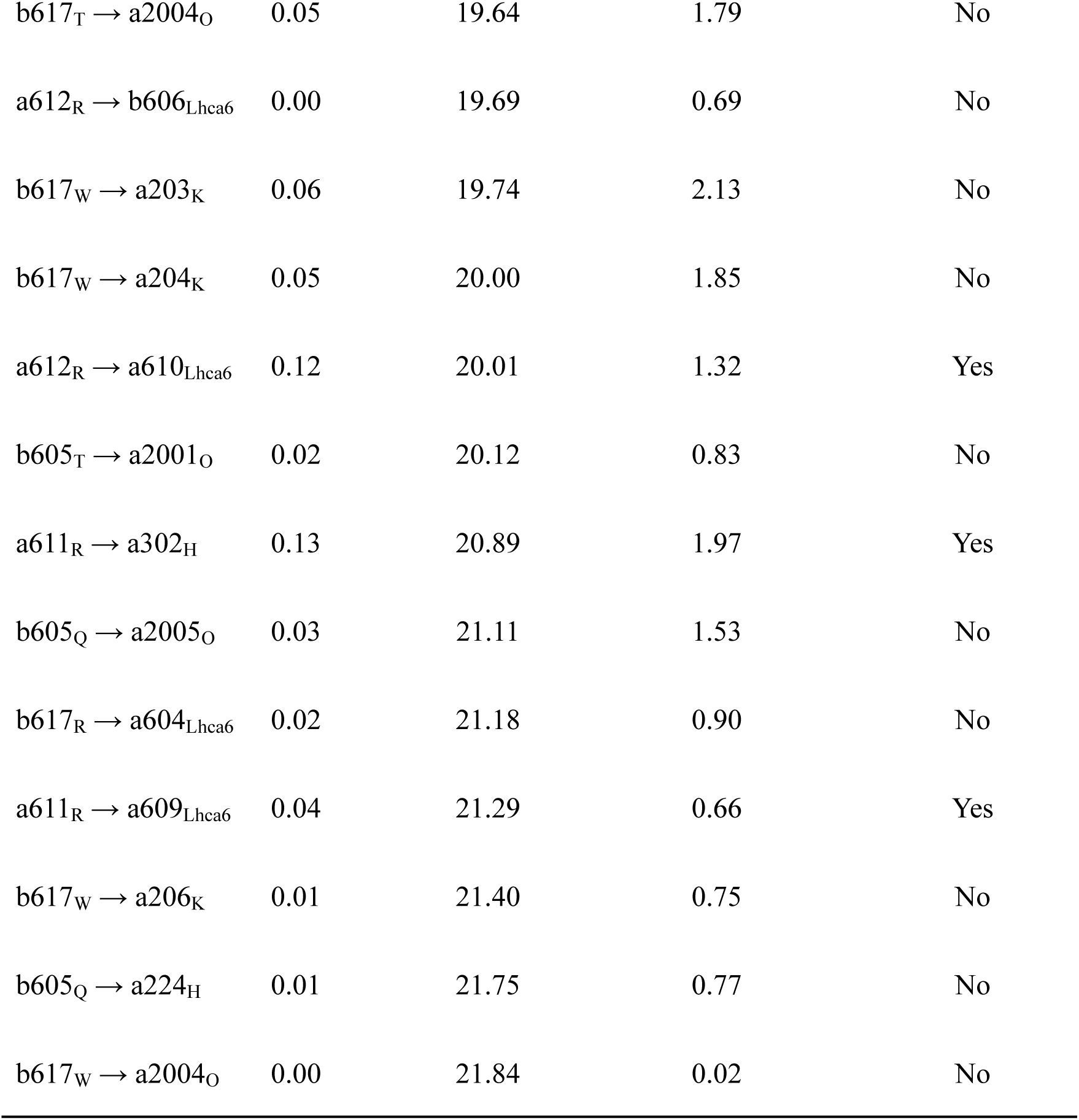
Calculated FRET rates between Lhcp and PSI core pigment pairs. In each row of the pigment pair column, on either side of the arrow, the first letter, the second value, and the third subscript represent the Chl species, the Chl number, and the polypeptide chain, respectively.

| Pigment pair | FRET rate | Dipole orientation |  | Chl *a* – Chl *a*<br>pair |
| --- | --- | --- | --- | --- |
| | $k_{\text{FRET}}$<br>(ps<sup>-1</sup>) | Distance | factor | |
| | | R (Å) | $K^2$ | |
| b605<sub>Q</sub> → a305<sub>L</sub> | 0.26 | 13.71 | 1.03 | No |
| b605<sub>W</sub> → a2004<sub>O</sub> | 0.22 | 14.67 | 1.33 | No |
| b608<sub>Q</sub> → a224<sub>H</sub> | 0.19 | 16.49 | 2.31 | No |
| a612<sub>W</sub> → a201<sub>K</sub> | 0.24 | 16.82 | 0.96 | Yes |
| a614<sub>R</sub> → b606<sub>Lhca6</sub> | 0.00 | 17.31 | 0.41 | No |
| b605<sub>T</sub> → a2003<sub>O</sub> | 0.03 | 17.52 | 0.61 | No |
| a610<sub>W</sub> → a201<sub>K</sub> | 0.00 | 19.16 | 0.00 | Yes |
| Mdp609<sub>Q</sub> → a224<sub>H</sub> | 0.05 | 19.26 | 1.57 | No |
| b606<sub>Q</sub> → a224<sub>H</sub> | 0.03 | 19.28 | 0.84 | No |
| $b617_T \rightarrow a2004_O$ | 0.05 | 19.64 | 1.79 | No |
| $a612_R \rightarrow b606_{Lhca6}$ | 0.00 | 19.69 | 0.69 | No |
| $b617_W \rightarrow a203_K$ | 0.06 | 19.74 | 2.13 | No |
| $b617_W \rightarrow a204_K$ | 0.05 | 20.00 | 1.85 | No |
| $a612_R \rightarrow a610_{Lhca6}$ | 0.12 | 20.01 | 1.32 | Yes |
| $b605_T \rightarrow a2001_O$ | 0.02 | 20.12 | 0.83 | No |
| $a611_R \rightarrow a302_H$ | 0.13 | 20.89 | 1.97 | Yes |
| $b605_Q \rightarrow a2005_O$ | 0.03 | 21.11 | 1.53 | No |
| $b617_R \rightarrow a604_{Lhca6}$ | 0.02 | 21.18 | 0.90 | No |
| $a611_R \rightarrow a609_{Lhca6}$ | 0.04 | 21.29 | 0.66 | Yes |
| $b617_W \rightarrow a206_K$ | 0.01 | 21.40 | 0.75 | No |
| $b605_Q \rightarrow a224_H$ | 0.01 | 21.75 | 0.77 | No |
| $b617_W \rightarrow a2004_O$ | 0.00 | 21.84 | 0.02 | No |

Fluorescence properties of A3 and A3L were characterized by fluorescence decay associated spectra at 77K (FDAS, Figure 1—supplement 2a-c). The fluorescence lifetimes of A3 were mainly made up of <10-ps and 65-ps components (Figure 1—supplement 2b), which correspond to the total trapping time around P700(Mimuro et al., 2010). The slight increase in the lifetimes of A3L compared to those of A3 (<10ps to 20ps, 65ps to 90ps) reflects the presence of larger antennae in A3L. The light-induced oxidation kinetics of P700 indeed showed that the PSI antenna size in A3L was clearly larger than that of A3 (Fig. 1c, Figure 1—supplement 3). Notably, the 77K fluorescence spectra of PSI–LHCI and PSI–LHCI–Lhcp exhibit a peak at 690 and 680 nm respectively (Fig. 1b), and there are no distinctive far-red peaks as reported previously(Swingley et al., 2010). The blue-shifted fluorescence was more prominent in PSI–LHCI–Lhcp fraction. These results and the EM analysis of the negatively stained particles (Figure 1—supplement 4 and Figure 1—table supplement 2) suggest that A3 almost exclusively consists of PSI–LHCI supercomplex, while A3L was mainly composed of PSI–LHCI–Lhcp supercomplex. We thus proceeded to solve the cryo-EM structure of the large PSI–LHCI–Lhcp supercomplex in A3L in order to reveal its detailed architecture.

### Supramolecular assembly of *Ot*Lhcp trimers (Trimers) with PSI–LHCI complex

Through the single-particle cryo-EM method, the structure of *Ot*PSI–LHCI–Lhcp supercomplex is solved at an overall resolution of 2.94 Å and the local regions of three Lhcp trimers (namely Trimers 1-3) are refined to 2.9-3.4 Å (Figure 2—supplement 1 and Figure 2—table supplement 1). As shown in Fig. 2, the *Ot*PSI–LHCI– Lhcp supercomplex consists of a central PSI monomer encircled by six LHCIs and three Trimers. *Ot*PSI contains two large core subunits (PsaA and PsaB), three subunits on the stromal surface (PsaC, PsaD and PsaE) (Fig. 2a-c), nine small membrane-embedded subunits (PsaF, PsaG, PsaH, PsaI, PsaJ, PsaK, PsaL, PsaM, PsaO), one subunit on the luminal surface (PsaN) (Fig. 2d-f), 314 Chls (245 Chl *a*, 60 Chl *b*, 9 Magnesium 2,4-divinylpheoporphyrin a_5_ monomethyl ester (Mdp), 106 carotenoids, two phylloquinones and three Fe_4_S_4_ clusters (Figure 2—table supplement 2). The densities for representative Chl, carotenoid and lipid molecules as well as two small subunits (PsaM and PsaN) are shown in Figure 2—supplement 2. To our best knowledge, the structure of *Ot*PSI has the largest number of subunits among the structures of PSI known so far (including those from plants, green algae, diatom, red algae, and cyanobacteria, Fig. 3).

**Fig. 2.**
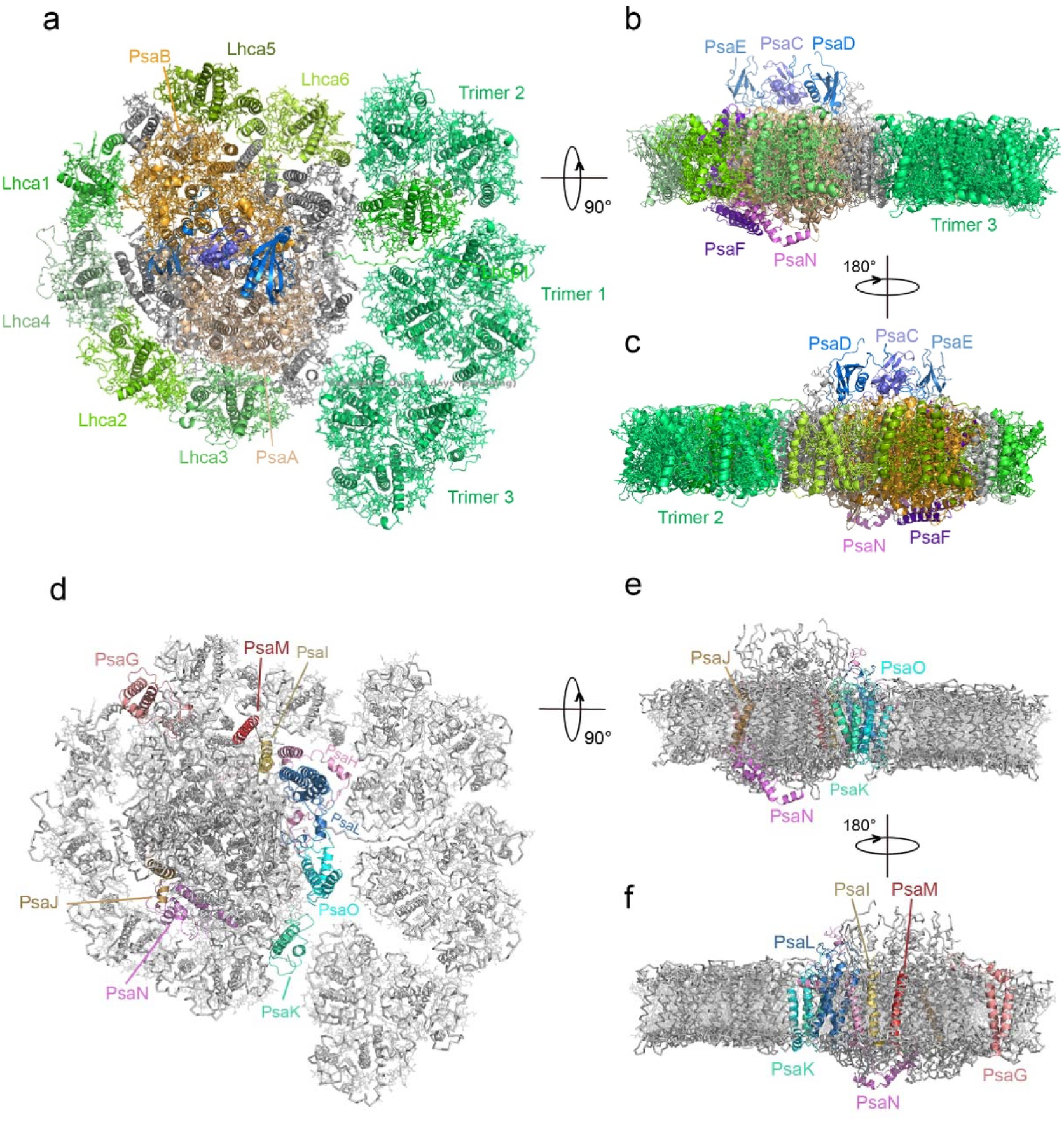
Overall architecture of *Ot*PSI–LHCI–Lhcp supercomplex. **a,** Top view of the supercomplex from stromal side along membrane normal. **b** and **c**, Two different side views of the supercomplex along the membrane plane. While the protein backbones are shown as cartoon models, the pigment and lipid molecules are presented as stick models. The iron-sulfur clusters are presented as sphere models. The bulk region of PSI core is colored in wheat, while Lhca1-6, Lhcp1 and Lhcp2, PsaC, PsaD, PsaE, PsaF and PsaN are highlighted in different colors. The remaining small subunits are in silver. **d-f**, The small subunits at the interfaces between PSI core and LHCI/Lhcp complexes. The viewing angles are the same as a-c, whereas the color codes are different. The interfacial small subunits are highlighted in various colors, while the PSI core and LHCI/Lhcp complexes are in silver. **Figure supplement 1.** Cryo-EM data collection, processing, refinement and validation statistics of OtPSI-LHCI-Lhcp structures. **Figure supplement 2.** Cryo-EM densities of various cofactors and protein subunits found in the PSI-LHCI-Lhcp supercomplex of *O. tauri* **Figure supplement 3.** The detailed local cryo-EM map features for identification of OtLhcp1 and OtLhcp2. **Table supplement 1.** Statistics of structural analysis of the OtPSI-LHCI-Lhcp supercomplex. **Table supplement 2.** Summarization of the components in the final structural model of the OtPSI-LHCI-Lhcp supercomplex.

**Fig. 3.**
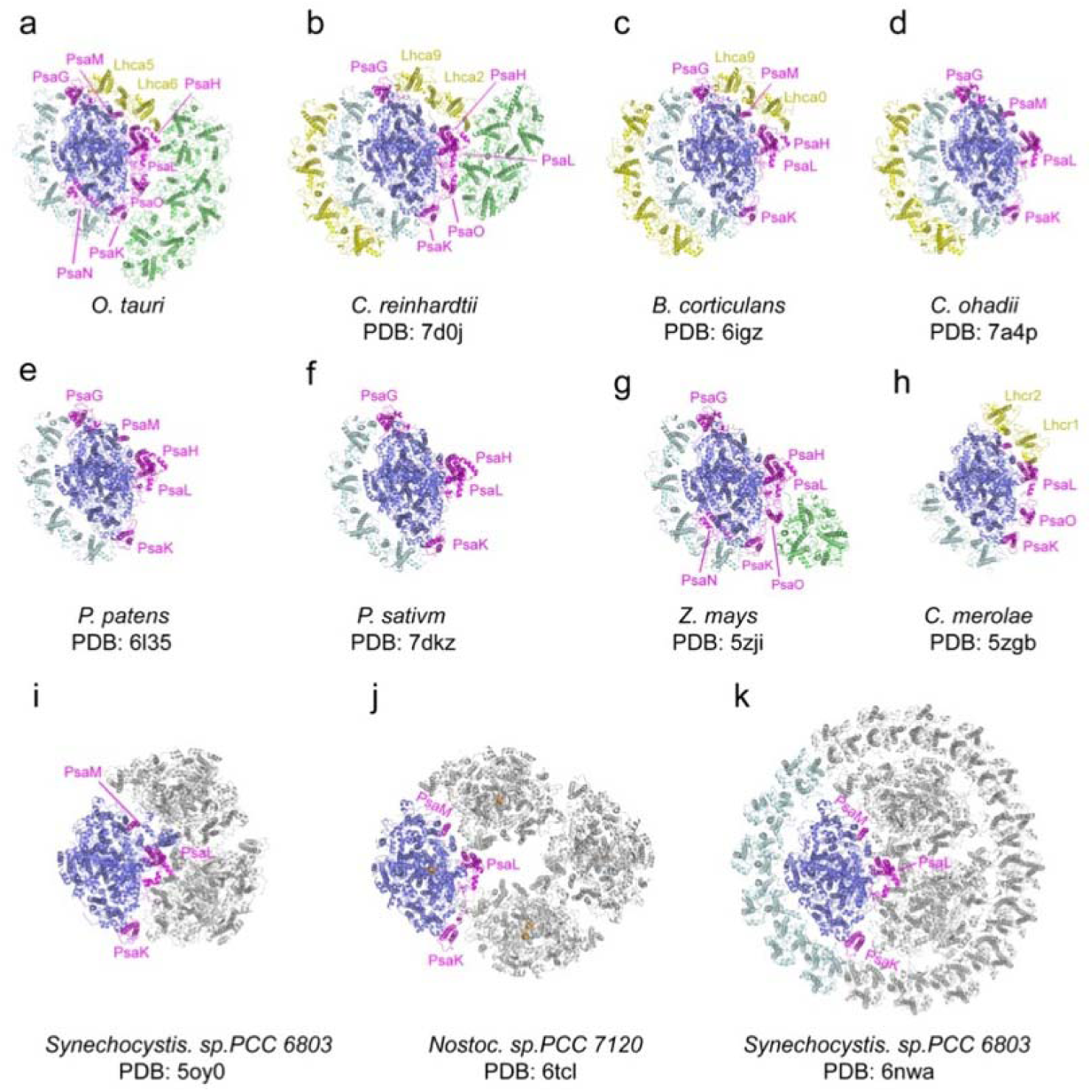
Comparison of *O. tauri* PSI–LHCI–Lhcp supercomplex with the PSI supercomplexes from other organisms. **a,** *Ot*PSI-LHCI-Lhcp supercomplex. **b**-**k,** Structures of PSI from difference species or at different states. Blue, large PSI core subunits; magenta, small PSI core subunits; yellow, Lhca5-Lhca6/Lhca2-Lhca9/Lhca0-Lhca9/Lhc1-Lhc2 and the outer belt of LHCI; cyan, Lhca1-Lhca2-Lhca3-Lhca4 and the inner belt of antenna complexes around PSI from other species; green, Lhcp or LHCII trimers; grey, symmetry-related units in trimeric or tetrameric PSI.

On one side of PSI, four Lhca (Lhca1-4) complexes fill in the gap between PsaG and PsaK (Fig. 2a), similar to those found in plant PSI–LHCI complexes(Mazor et al., 2017; Qin et al., 2015) (Fig. 3). On the other side, a Lhca5-Lhca6 heterodimer associate with PSI through PsaG, PsaB, PsaI, PsaM and PsaH subunits (Fig. 2a and d). The location of Lhca5-Lhca6 heterodimer overlaps with that of Lhca2-Lhca9 or Lhcr1-Lhcr2 heterodimer associated with *C. reinhardtii*(Su et al., 2019; Suga et al., 2019) or *C. merolae* PSI(Pi et al., 2018), whereas the corresponding sites are vacant in plant PSI structures(Mazor et al., 2017; Pan et al., 2018; Qin et al., 2015; Yan et al., 2021). While the fourth transmembrane helix of *Cr*Lhca2 or Lhca5 from *Dunaliella salina* (also known as transmembrane helix F/TMF or TMH4), located at the dimerization interface of the Lhca2/Lhca9 or Lhca5-Lhca6 heterodimer, was previously proposed to replace the role of PsaM in mediating assembly of the LHCI heterodimer with PSI(Caspy et al., 2020; Suga et al., 2019), the fourth helix of *Ot*Lhca6 and PsaM coexist in *O. tauri*. As PsaM is also present in moss(Gorski et al., 2022) and cyanobacteria(Jordan et al., 2001), where LHCI heterodimer is absent at this position, the role and the origin of the fourth helix of the Lhca proteins might not be directly related to PsaM and need be revisited.

Previously, it was found that plant and *C. reinhardtii* PSI complexes contain some red-form Chls, absorbing photons at energy levels below that of the primary donor and mainly located in the LHCI complexes(Croce & van Amerongen, 2013). Unlike plant and *C. reinhardtii* PSI, *Ot*PSI–LHCI–Lhcp supercomplex sample does not exhibit far-red peaks in the 77-K fluorescence spectra (Fig. 1b). In the *Ot*PSI–LHCI, the Chl *a*603-*a*609 pairs of Lhca1-4 (corresponding to the red-form Chls found in plant and *C. reinhardtii* Lhca1-4) are separated at larger distances, while those in Lhca5 and Lhca6 are similar to the ones in Lhca9 and Lhca2 from *C. reinhardtii* (Fig. 4). Moreover, the axial ligands of Chl *a*603 in Lhca1-4 from *O. tauri* are all His residues instead of Asn (Fig. 4b-e). Mutation of Asn to His for the ligand of Chl *a*603 in plant Lhca3 and Lhca4 led to absence of red spectral forms, while substitution of Asn for His in Lhca1 caused red shift of the fluorescence emission(Morosinotto et al., 2003). As His has longer side chain than Asn, the position of Chl *a*603 in *Ot*Lhca1-4 is located farther away from the protein backbone (in comparison with those from plant and *C. reinhardtii* LHCIs) so that the distance between Chl *a*603 and Chl *a*609 becomes larger and their excitonic coupling strength might be reduced as a result. The axial ligands of Chl *a*603 in Lhca5-Lhca6 dimer from *O. tauri* are both Asn residues same as those in the Lhca9-Lhca2 dimer on a similar location in *C. reinhardtii* (Fig. 4b-e). Although Chl *a*603-*a*609 pairs of Lhca2 and Lhca9 were proposed to be responsible for the red spectral forms in *C. reinhardtii*(Mozzo et al., 2010), those in *Ot*Lhca5 and *Ot*Lhca6 with similar configuration do not cause the red spectral forms. This might be due to their different local environments.

**Fig. 4.**
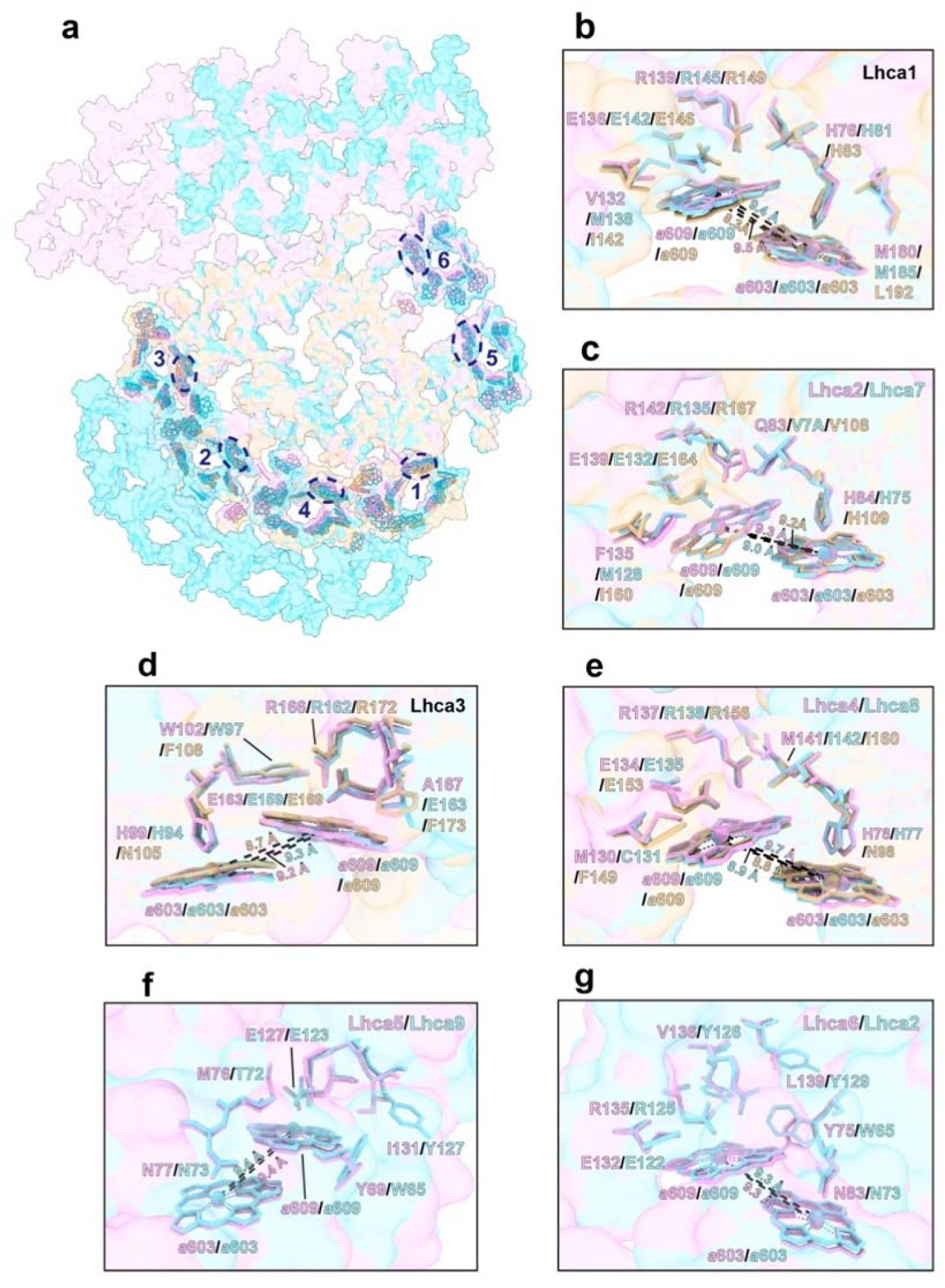
Comparison of Chl *a*603–*a*609 dimer in Lhca complex among *O. tauri*, *C. reinhardtii* and *P. sativum*. **a**, Overall arrangement of the Chl *a*603-*a*609 dimers in six Lhca complexes of *O. tauri*, *C. reinhardtii* and *P. sativum*. The chlorophylls in the PSI supercomplexes are presented in surface models and those of Lhca complexes are superposed with stick models. Color code : pink, *O. tauri*; golden, *P. sativum*; light blue, *C. reinhardtii*. The blue labels (1-6) indicate the Lhca1-Lhca6 subunits and the dashed ovals label the locations of the *a*603-*a*609 dimer in each LHCI. **b-g**, Comparison of the Chl *a*603-*a*609 dimers and their local environments in six different Lhca subunits from the three different species. Note that the axial ligands of Chl a603 in Lhca1-4 from *O. tauri* are all His, while the Asn in the 603 site of plant Lhca3 and Lhca4 are crucial for the formation of the red-most form chlorophyll. The Lhca5/Lhca9 and Lhca6/Lhca2 are only present in *O. tauri* and *C. reinhardtii* but absent in *P. sativum.* The *a*603-*a*609 dimer and the key amino acid residues around the two chlrophylls are highlighted as stick models. The number labeled nearby the black dashed lines indicate the Mg-Mg distances between Chl *a*603-*a*609 dimer in Lhca complexes from *O. tauri*, *C. reinhardtii* and *P. sativum*.

Besides the six LHCI complexes, three Trimers 1-3 binds to the PSI–LHCI complex on the third side between Lhca6 and PsaK (Fig. 2). As a result, the PSI core is enclosed by an irregular annular belt formed by the LHCI, Lhcp complexes, PsaG and PsaK (Fig. 2a and d). This side of the PSI core was partly filled by one LHC trimer or two LHC trimers in higher plants or green algae when they are under state 2 conditions(Huang et al., 2021; Pan et al., 2018; Pan et al., 2021).

Among the three trimers associated with *Ot*PSI, Trimer 1 and 3 are both (Lhcp2)_3_ homotrimers, whereas Trimer 2 is a Lhcp1(Lhcp2)_2_ heterotrimer. The detailed cryo-EM map features for identification of Lhcp1 and Lhcp2 are shown in Figure 2—supplement 3. Trimer 1 and Trimer 3 bind to PSI on the PsaO and PsaK sides respectively, while Trimer 2 assembles with PSI on the PsaL-PsaH side through Lhcp1 subunit and interacts with Lhca6 through a Lhcp2 subunit. As Trimer 1 is sandwiched between Trimers 2 and 3, it forms close contacts with both Trimer 2 and 3, and is related with them through pseudo-C2 symmetry axes at their interfaces. In *C. reinhardtii*, two LHCII trimers associate with PSI in State 2(Huang et al., 2021; Pan et al., 2021), whereas one LHCII trimer is located at the peripheral region of *Zea mays* PSI (*Zm*PSI) in State 2(Pan et al., 2018) (Fig. 3). While the binding sites of Trimer 1 and 2 partially overlaps with those of LHCII-1 and LHCII-2 trimers from *Cr*PSI–LHCI-LHCII supercomplexes respectively, they do not superpose well with each other (Fig. 5). The LHCII trimer associated with *Zm*PSI binds to a position between Lhcp timers 1 and 3.

**Fig. 5.**
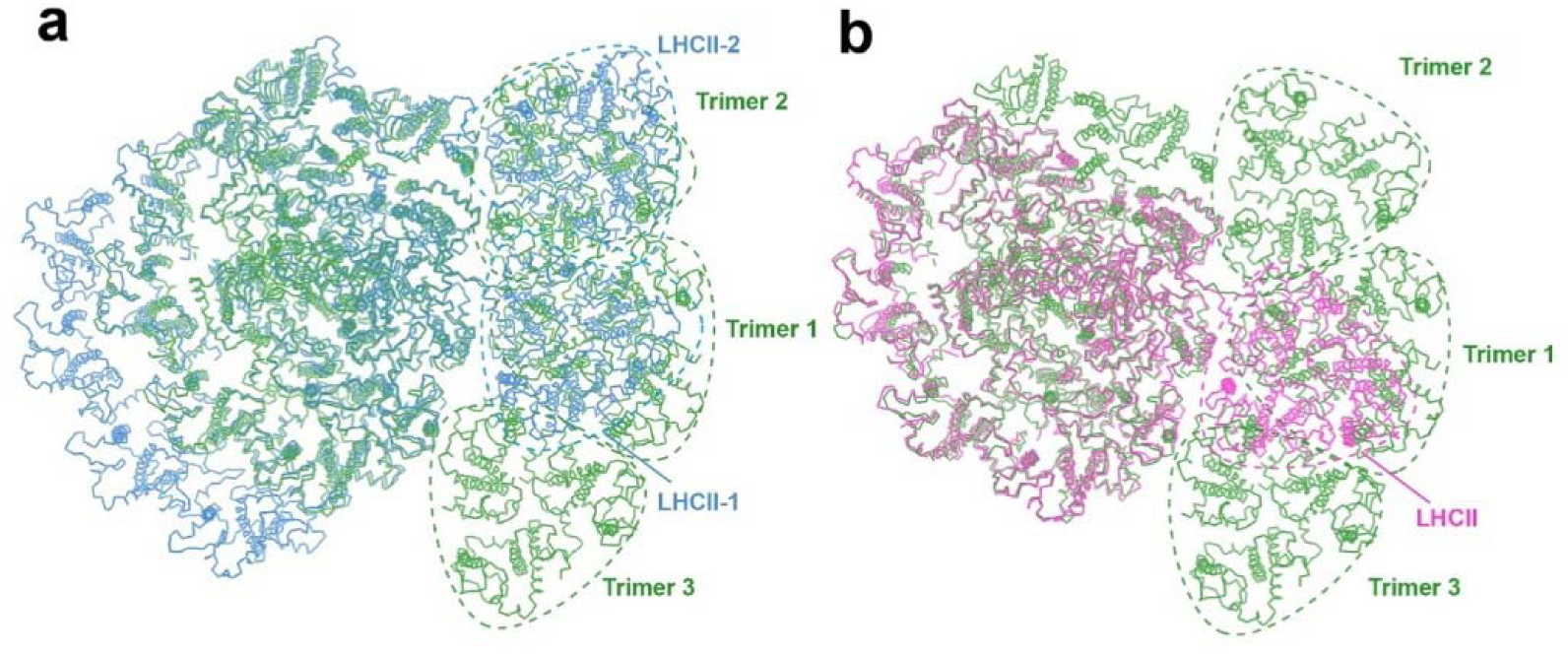
Comparing the binding sites of Trimers with those of LHCII trimers bound to *C. reinhardtii* and plant PSI. **a and b**, The structure of *Ot*PSI-LHCI-Lhcp supercomplex is superposed with the PSI-LHCI-LHCII supercomplex from *C. reinhardtii* (PDB code: 7DZ7, a) or *Z. mays* (PDB code: 5ZJI, b). The three structures are superposed on the common PsaA subunits. The dash triangular rings outline the approximate boundaries of Trimers or LHCII trimers. Color code: green, *Ot*PSI-LHCI-Lhcp; blue, *Cr*PSI-LHCI-LHCII; magenta, *Zm*PSI-LHCI-LHCII.

Trimer 1 is located nearby PsaO and the closest interfacial distances between them are 7.1 Å or larger (Fig. 6a). Although Trimer 1 does not form direct interactions with PsaO, there might be some unobserved lipid molecules at the interface to mediate their assembly. On the other hand, Trimer 2 and Trimer 3 form close and direct interactions with PsaL-PsaH and PsaK, respectively (Fig. 6b and c). Trimer 2 binds to PSI on three different sites (Fig. 6d-f). At site 1, Lhcp1_Trimer2_ has its elongated N-terminal region (NTR) partially inserted into a surface pocket formed by PsaL and PsaH subunits (Fig. 6b). The NTR of Lhcp1 contains an RRpT (pT, phosphorylated Thr residue) motif identical to those found in *Z. mays* pLhcb2(Pan et al., 2018) and *C. reinhardtii* pLhcbM1(Pan et al., 2021). The RRpT motif of Lhcp1 interacts with nearby amino acid residues through salt bridges and hydrogen bonds (Fig. 6d), in a similar way as those of *Z. mays* pLhcb2 and *C. reinhardtii* pLhcbM1. Sites 2 and 3 are located in the membrane-embedded regions on the stromal and luminal sides, respectively (Fig. 6b). On these two sites, Lhcp1 forms hydrogen bonds and van der Waals interactions with nearby amino acid residues and pigment molecules from PsaH and PsaL subunits (Fig. 6e and f). Meanwhile, Trimer 3 associates with PsaK mainly through van der Waal interactions on the stromal and luminal sides (Fig. 6g).

**Fig. 6.**
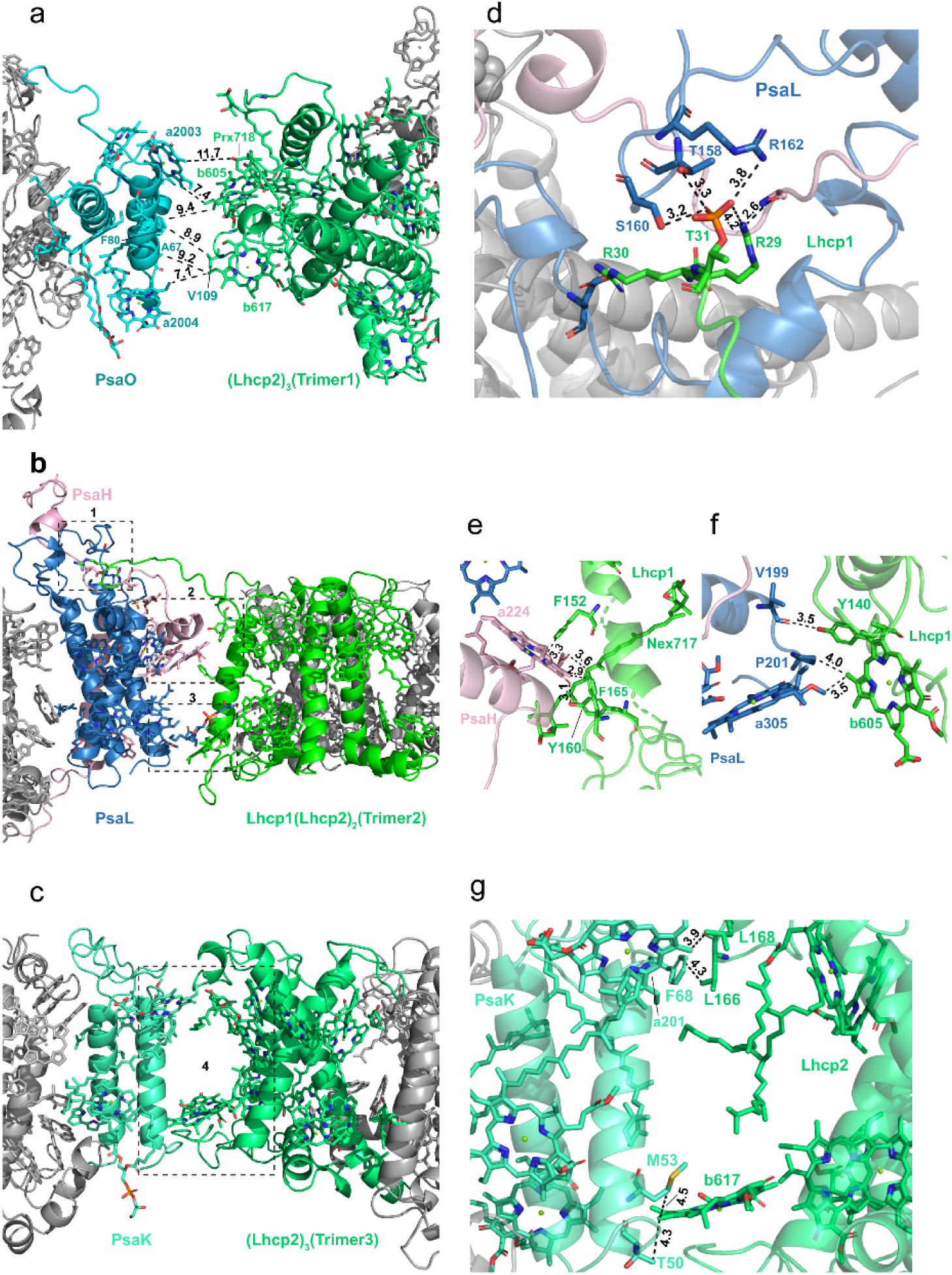
Interfaces between Trimers and PSI subunits. **a,** The interface between Trimer 1 and PsaO. **b,** The interface between Trimer 2 and PsaL-PsaH. **c**, The interface between Trimer 3 and PsaK. **d-f**, The detailed interactions between Lhcp1 of Trimer 2 and PsaL/PsaH at sites 1-3 shown in **b**. **g**, The detailed interactions between Lhcp2 of Trimer 3 and PsaK at site 4 shown in **c**. The numbers labeled nearby the dash lines are distances (Å) between two adjacent groups.

### Characteristic features of Lhcp1 and Lhcp2 monomers

As an ancient member of the green lineage, *O. tauri* belongs to Mamiellales of Prasinophyceae at the basal position of the green lineage. While *Ot*Lhcp1 and *Ot*Lhcp2 are evolutionarily related to plant Lhcbs and *Cr*LhcbMs(Six et al., 2005), they only share low sequence identity, e.g. 32-37% sequence identity with *Zm*Lhcb2 or *Cr*LhcbM1. The apoproteins of *Ot*Lhcp1 and *Ot*Lhcp2 adopt a classical fold of Lhc family with three transmembrane helices (A, B and C) and three short amphipathic helices on the luminal side (D, E and F) (Fig. 7a, b and Figure 7—supplement 1). While the structure of *Ot*Lhcp1 highly resembles that of *Ot*Lhcp2, it differs from those of *Zm*Lhcb2 and *Cr*LhcbM1 in the N-terminal region, EC loop, AC loop and CTR (Figure 7—supplement 2a-c). Besides, helices B, A and C of *Ot*Lhcp1/2 are slightly shorter than the corresponding ones in *Zm*Lhcb2 or *Cr*LhcbM1 (Figure 7—supplement 2b and c).

**Fig. 7.**
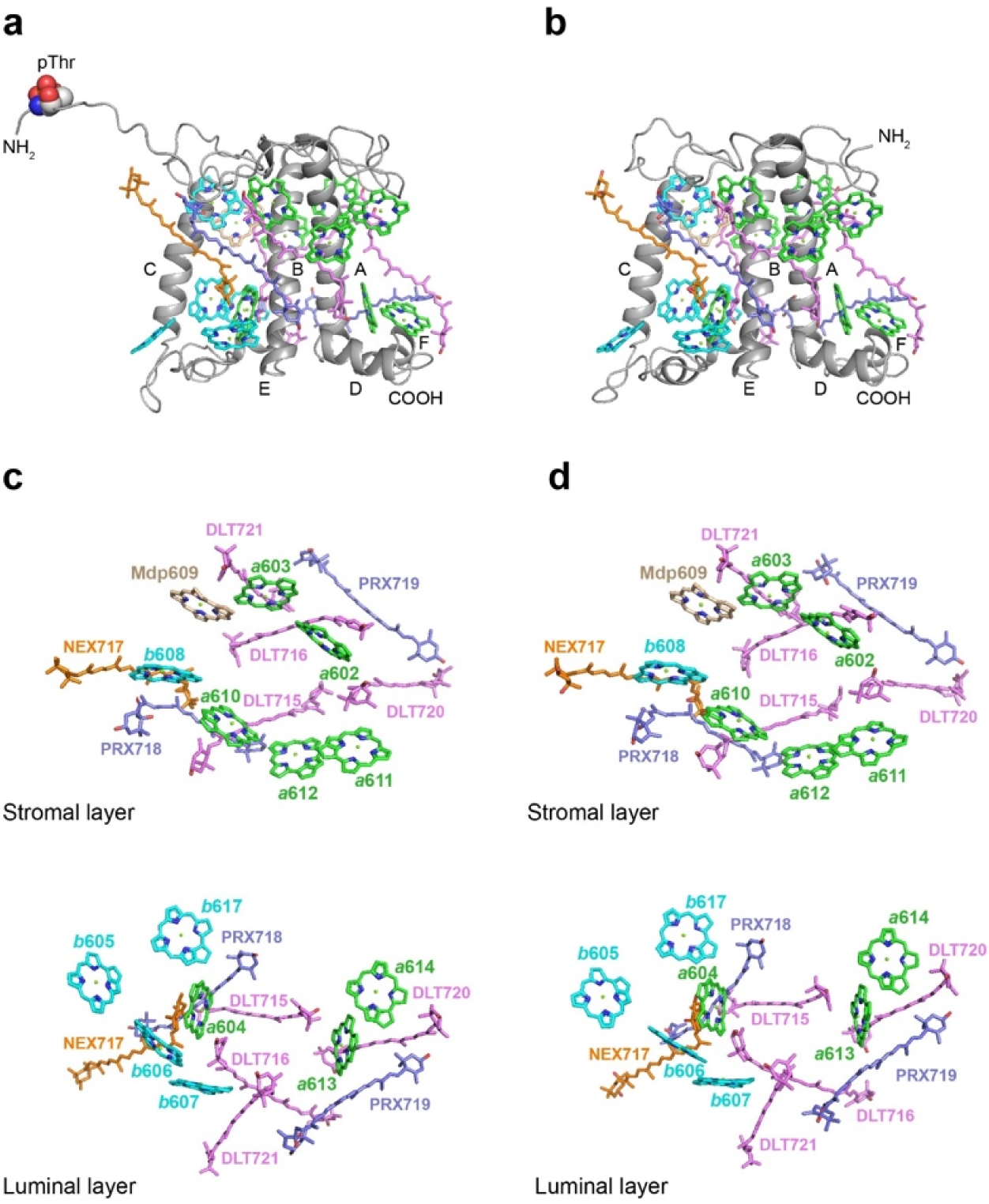
The structures and pigment compositions of *Ot*Lhcp1 and *Ot*Lhcp2. ***a* and *b***, Side views of *Ot*Lhcp1 (a) and *Ot*Lhcp2 (b) structures. The backbones of *Ot*Lhcp1/*Ot*Lhcp2 apoproteins are showed as cartoon models, while the pigments are presented as stick models. The phytyl chains of chlorophyll molecules are omitted for clarity. The phosphorylated Thr residue at the N-terminal region of Lhcp1 is highlighted as sphere models. A, B and C indicate the three transmembrane helices in *Ot*Lhcp1/*Ot*Lhcp2 apoproteins, whereas D and E are the two amphipathic helices at the luminal surface. Color codes: gray, Lhcp1 and Lhcp2 apoproteins; green, Chl *a*; cyan, Chl *b*; magenta, dihydrolutein/DLT; purple, prasinoxanthin/PRX; orange, neoxanthin/NEX. **c and d**, The arrangement of pigment molecules in *Ot*Lhcp1 (c) and *Ot*Lhcp2 (d). For clarity, the apoproteins are not shown. In the upper row, only pigment molecules within the layer close to stromal surface are shown, while the lower row shows the pigment molecules within the layer close to the luminal surface. **Figure supplement 1.** Cryo-EM densities of OtLhcp1 and OtLhcp2. **Figure supplement 2.** Superposition of the *Ot*Lhcp1 structure with those of *Ot*Lhcp2, *Zm*Lhcb1 and *Cr*LhcbM1.

In terms of pigment composition, *Ot*Lhcp1 and *Ot*Lhcp2 each contain 14 Chl molecules (8 Chl *a*, 5 Chl *b* and 1 Mdp) and 7 carotenoid molecules (4 Dlt, 2 Prx and 1 Nex) (Fig. 7a and b). The Chl *b*/*a* ratio of the structural model is 0.625, close to the previously reported value of 0.736 for the Lhcp2 preparation(Swingley et al., 2010). While the number of Prx molecules found in the structure matches the previous prediction, four (instead of one) Dlt molecules are more than expected(Swingley et al., 2010). As *O. tauri* OTH95 species thrives in lagoons and shallow area of the ocean frequently challenged by high light with an intensity up to 2,000 μmol photons m^-2^sec^-1^ (Courties et al., 1994), incorporation of a large number of carotenoid molecules in *Ot*Lhcp1 and *Ot*Lhcp2 may help to protect the algae from the damaging effect of harmful excess excitation energy, by quenching triplet-state Chl, scavenging toxic oxygen species or quenching of the singlet excited-state Chl(Bassi & Dall’Osto, 2021; Young, 1991).

The Chl molecules in *Ot*Lhcp1/*Ot*Lhcp2 are arranged into two membrane-embedded layers close to the stromal and luminal surfaces, respectively (Fig. 7c and d). The stromal and luminal layer each has 7 Chl molecules. While the stromal layer (5 Chl *a*, 1 Chl *b* and 1 Mdp) contains more Chl *a* than Chl *b*, the luminal layer (3 Chl *a* and 4 Chl *b*) harbors more Chl *b* than Chl *a*. In the stromal layer, six Chls (*a*602, *a*603, *b*608, *a*610, *a*611 and *a*612) can find their counterparts of the same identity in plant or *C. reinhardtii* LHCII (Figure 7—supplement 2d-f). The 609 site is occupied by an Mdp molecule in *Ot*Lhcp1/2, whereas LHCII has the 609 site occupied by a Chl *b*(Liu et al., 2004). The Chl *b*609 in LHCII has its C7-formyl group forming a hydrogen bond with Gln131 on the Lhcb1 protein, and the side chain amine group of Gln131 is crucial for selective binding of Chl *b*609 and *b*607 in LHCII(Bassi et al., 1999). Lhcp1 and Lhcp2 adopt a Glu residue in the position corresponding to Gln131 of the Lhcb1 protein (Fig. 8). Similar change is also observed in CP29 or CP26 with lower Chl *b* content than LHCII, and Q131E mutation of Lhcb1 led to decrease of Chl *b* content and increase of Chl *a* content(Bassi et al., 1999). As the side chain hydroxyl group of Glu150/Glu129 residue in *Ot*Lhcp1/2 forms a hydrogen bond with Chl *b*607, the 609 site is occupied by an Mdp molecule with a C^7^-methyl group (instead of a Chl *b*) due to lack of hydrogen bond donor. Moreover, Chl *b*601 found in plant and *C. reinhardtii* LHCII is absent in *Ot*Lhcp1/2, mainly because the head group of a carotenoid molecule (Dlt720) occupies part of its binding site (Figure 7—supplement 2e and f) and the NTR adopts a conformation different from those of LHCII apoproteins Figure 7—supplement 2b and c).

**Fig. 8.**
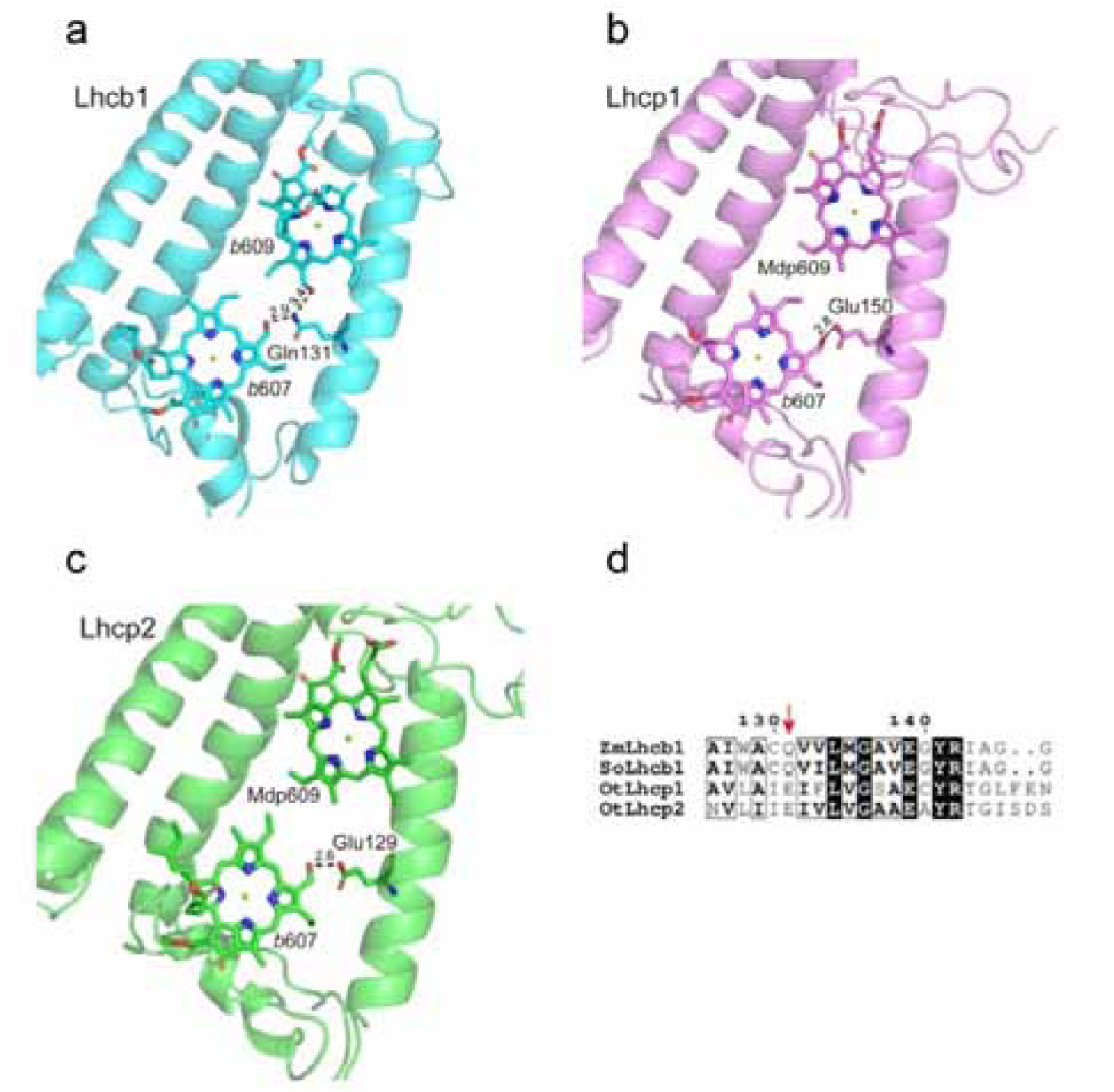
The key amino acid residues related to the selectivity of chlorophyll binding site 609 in *Ot*Lhcp1/2 and plant Lhcb1. **a,** The side chain amine group of Gln131 in Lhcb1 from *Spinacea oleracea* (*So*Lhcb1) offers two hydrogen-bond donors for selective binding of two Chl *b* molecules. **b and c**, The carboxyl group of Glu150/Glu129 in *Ot*Lhcp1/2 provides only one hydrogen-bond donor for selective binding of only one Chl *b* molecule instead of two. **d**, Sequence alignment analysis of the plants Lhcb1 and *Ot*Lhcp1/2. Color codes: cyan, Lhcb1; violet, Lhcp1; green, Lhcp2. The dash lines indicate the putative hydrogen bonds between the C7-formyl groups of Chl *b* molecules and Glu/Gln residues. The numbers labeled nearby the dash lines are the distances (Å) between hydrogen bond donor and acceptors. The pigment molecules are shown as stick models, while the protein backbones are presented as cartoon models. **Figure supplement 1.** Sequence alignment of OtLhcp1 and OtLhcp2 with LhcbM1 from *C. reinhardtii* and Lhcb2 from *Z. mays*. **Figure supplement 2.** The detailed features at the trimerzation interfaces of OtTrimers

In the luminal layer, six of the seven Chls (*a*604, *b*605, *b*606, *b*607, *a*613 and *a*614) can find their counterparts in plant LHCII. A new Chl (Chl *b*617, absent in plant or *C. reinhardtii* LHCII) is found in the EC-loop region of *Ot*Lhcp1/2 (Fig. 7c and d, Figure 7—supplement 2d) and is likely coordinated by the backbone carbonyl of a proline residue conserved in both Lhcp1 and Lhcp2, but not in plant Lhcb2 or *Cr*LhcbM1 (Figure 8—supplement 1). The binding site of Chl *b*617 is located in a motif of the EC loop region different from the corresponding region in plant Lhcb2 or *Cr*LhcbM1. Among the 7 carotenoid molecules, six of them span the membrane and connect the luminal-layer Chls with those in the stromal layer, whereas one (Prx719) lies horizontally in the luminal layer and contributes to trimerization of *Ot*Lhcp1 and *Ot*Lhcp2 (Figure 8—supplement 2). While three carotenoids (Dlt715, Dlt716 and Nex717) of *Ot*Lhcp1/2 can found their counterparts in *Zm*Lhcb2 or *Cr*LhcbM1, the other four (Prx718, Prx719, Dlt720 and Dlt721) are specific to *Ot*Lhcp1/2 and absent in *Zm*Lhcb2 or *Cr*LhcbM1 (Figure 7—supplement 2e and f). Moreover, the violaxanthin molecule found at the trimerization interface of plant LHCII(Liu et al., 2004) is absent in *Ot*Lhcp1/2.

### Formation of Lhcp1–(Lhcp2)_2_ and (Lhcp2)_3_ trimers

Previously, it was suggested that Lhcp2 from *O. tauri* may form a hexameric unit according to the gel filtration analysis result(Swingley et al., 2010). There are one Lhcp1 and eight Lhcp2 associated with *Ot*PSI–LHCI, forming three individual trimers (Trimers 1-3, Fig. 2a). Among them, Trimer 1 and Trimer 2 interact with each other closely and may form a stable hexameric unit. Despite that Trimer 2 differs from the other two trimers in terms of subunit composition [Lhcp1(Lhcp2)_2_ vs (Lhcp2)_3_], the overall structures of the three trimers are highly similar (Fig. 9a). The NTR, helix C, helix E, the E-C loop and the C-terminal region (CTR) of Lhcp1 and Lhcp2 are located at the interfaces between adjacent monomers (Fig. 9a and b). Three carotenoids, namely Prx719, Dlt720 and Dlt721, intercalate into the space between adjacent monomers and interact closely with nearby cofactors and amino acid residues (Fig. 9b). Prx719 spans a Trimer by extending from the periphery to the center on the luminal side and forms hydrogen bonds and van der Waals interactions with nearby CTR amino acid residues (FKFY/FGFY motif, Figure 8—supplement 1, Figure 8—supplement 2e-f). Meanwhile, Dlt720 and Dlt721 are mainly located at the peripheral and middle regions of the trimerization interface respectively (Fig. 9e). Thereby, the interfacial carotenoids serve as molecular staples, holding the three monomers together and stabilizing the trimeric assembly of Lhcps.

**Fig. 9.**
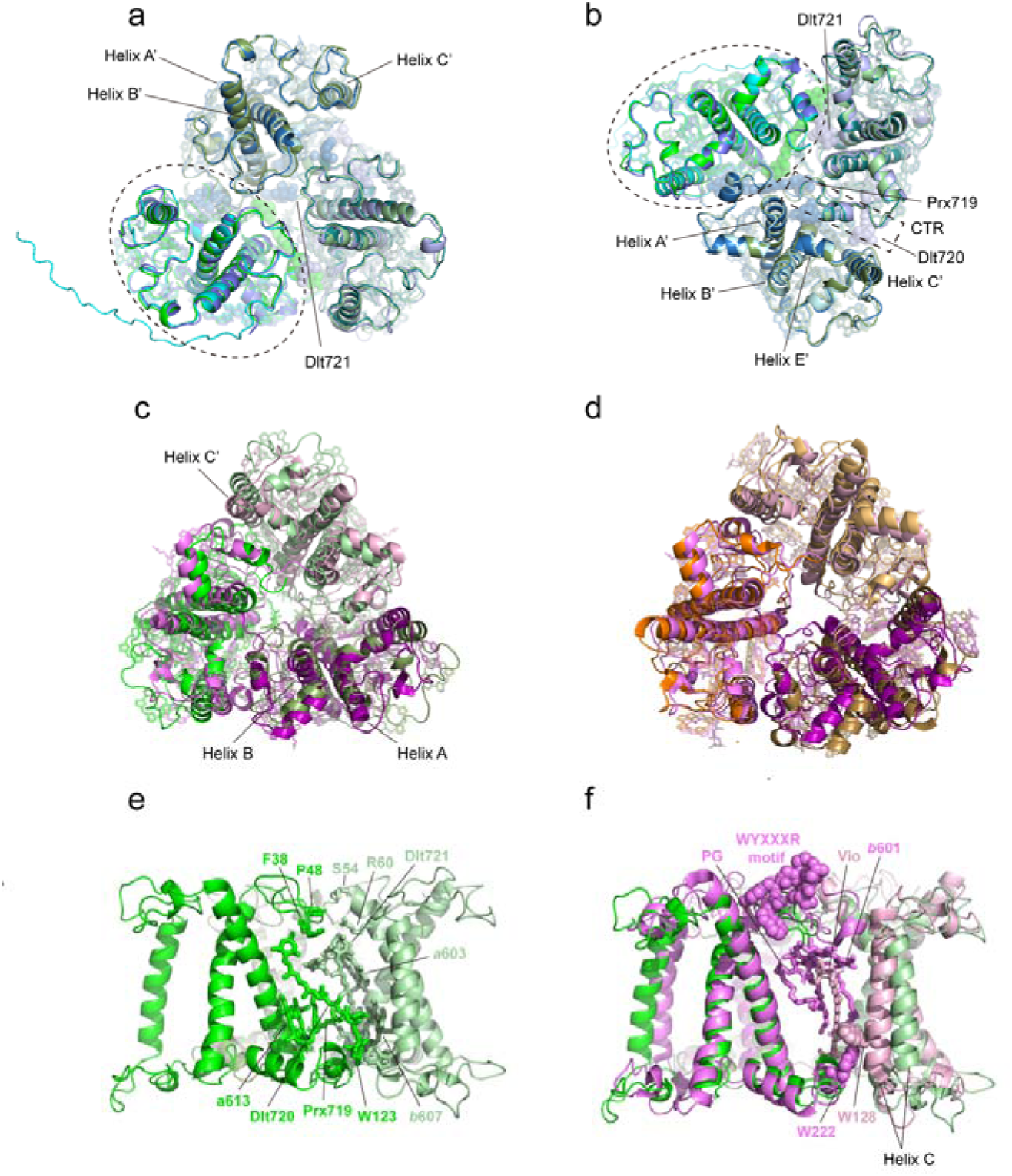
Monomer–monomer interface of the Trimers. **a and b,** Superposition of the three Trimers associated with PSI viewed from stromal (a) and luminal (b) sides. Color codes: green/pale green/dark green, three different monomers of Trimer 1 [(Lhcp2)_3_]; cyan/pale cyan/dark cyan, three different monomers of Trimer 2 [Lhcp1-(Lhcp2)_2_]; blue/light blue/dark blue, three different monomers of Trimer 3 [(Lhcp2)_3_]. The dashed elliptical ring indicates the approximate boundary of one Lhcp1/2 monomer. The dashed box outlines the interfacial location of the C-terminal regions (CTR) of Lhcp1/2. The three carotenoid molecules located at the monomer-monomer interfaces are highlighted as sphere models, while the remaining cofactors are shown as stick models. **c**, The (Lhcp2)_3_ trimer superposed on spinach LHCII trimer. The three monomers of spinach LHCII trimer (PDB code: 1RWT) are colored in violet, light pink and deep purple respectively, whereas Lhcp2 monomers are in green. **d**, Superposition of spinach LHCII trimer with *C. reinhardtii* LHCII trimer. The three monomers of *C. reinhardtii* LHCII (PDB code: 7D0J) are colored in orange, light orange and sand. **e,** Side view of the monomer-monomer interface of the (Lhcp2)_3_ trimer. The amino acid residues and cofactors in contact with the adjacent monomer are highlighted as sticks models. **f**, The monomer-monomer interface of (Lhcp2)_3_ trimer from *O. tauri* compared to the one in spinach LHCII trimer. The cofactors and amino acid residues involved in trimerization of spinach LHCII trimer, but absent in *Ot*(Lhcp2)_3_ trimer, are shown as sphere models. NTR, the N-terminal region; CTR, the C-terminal region.

Like the *Ot*Trimers, the major LHCII from plants and *C. reinhardtii* also exists in homotrimeric or heterotrimeric states(Caffarri et al., 2004; Kawakami et al., 2019; Standfuss & Kühlbrandt, 2004). Trimerization of LHCII is crucial for thermal stability of the complex, may modulate the pigment domains related to nonphotochemical quenching of Chl fluorescence and allows the transfer of excitation energy between adjacent monomers through interfacial Chls(Novoderezhkin et al., 2011; Wentworth et al., 2004). While the *Ot*Lhcp2 and LHCII trimers share similar overall shapes, they differ from each other in terms of mutual positions and distances between adjacent monomers (Fig. 9c and d), and the interfacial cofactors and amino acid residues (Fig. 9e and f). Besides, the phosphatidylglycerol (PG) molecule, Chl *b*601, Vio, the WYXXXR motif (also known as trimerization motif) at the NTR(Koziol et al., 2007) and two bulky amino acid residues (Trp128 and Trp222) found at the trimerization interface of plant LHCII trimer are all absent in the *Ot*Trimers (Figure 8—supplement 1).

### Potential energy transfer pathways between Trimers and PSI

Among the three Trimers associated with PSI, Trimer 1 may have a crucial role in mediating the excitation energy transfer between Trimer 2 and Trimer 3. As shown in Fig. 10, the Chl molecules in Trimer 1 are well connected with those from Trimer 2 and Trimer 3 through multiple pairs of interfacial Chls in the stromal and luminal layers. Remarkably, the distances between adjacent Chls at the trimer-trimer interface are comparable to or even smaller than those of inter-monomer Chl pairs within the individual trimers (Fig. 10a-d). In the stromal layer, the Chl *a*611-*a*612 dimer from monomer S_Trimer1_ is tightly connected with the Chl *a*611-*a*612 dimer from monomer Q_trimer2_ at 12.3 or 12.6 Å distances (Mg-Mg, Fig. 10a). Meanwhile, Chl *a*611 from monomer T_Trimer2_ is connected with Chl *a*611 from monomer V_Trimer3_ of at 17.9 Å distance. In the luminal layer of the trimer, Chl *a*614 and *b*617 from monomer S_Trimer1_ are connected with Chl *b*617 and *a*614 from monomer Q_Trimer2_, respectively (Fig. 10b). Meanwhile, Chl *a*614 from monomer T_Trimer1_ and Chl *b*605 from monomer U_Trimer1_ make very close connections with Chl *a*614 and *b*617 from monomer V_Trimer3_ at 14.7 and 9.6 Å distances, respectively. The close relationships among the three Trimers may allow the excitation energy to be shared efficiently among them through the closely-connected Chl pairs at their interfaces.

**Fig. 10.**
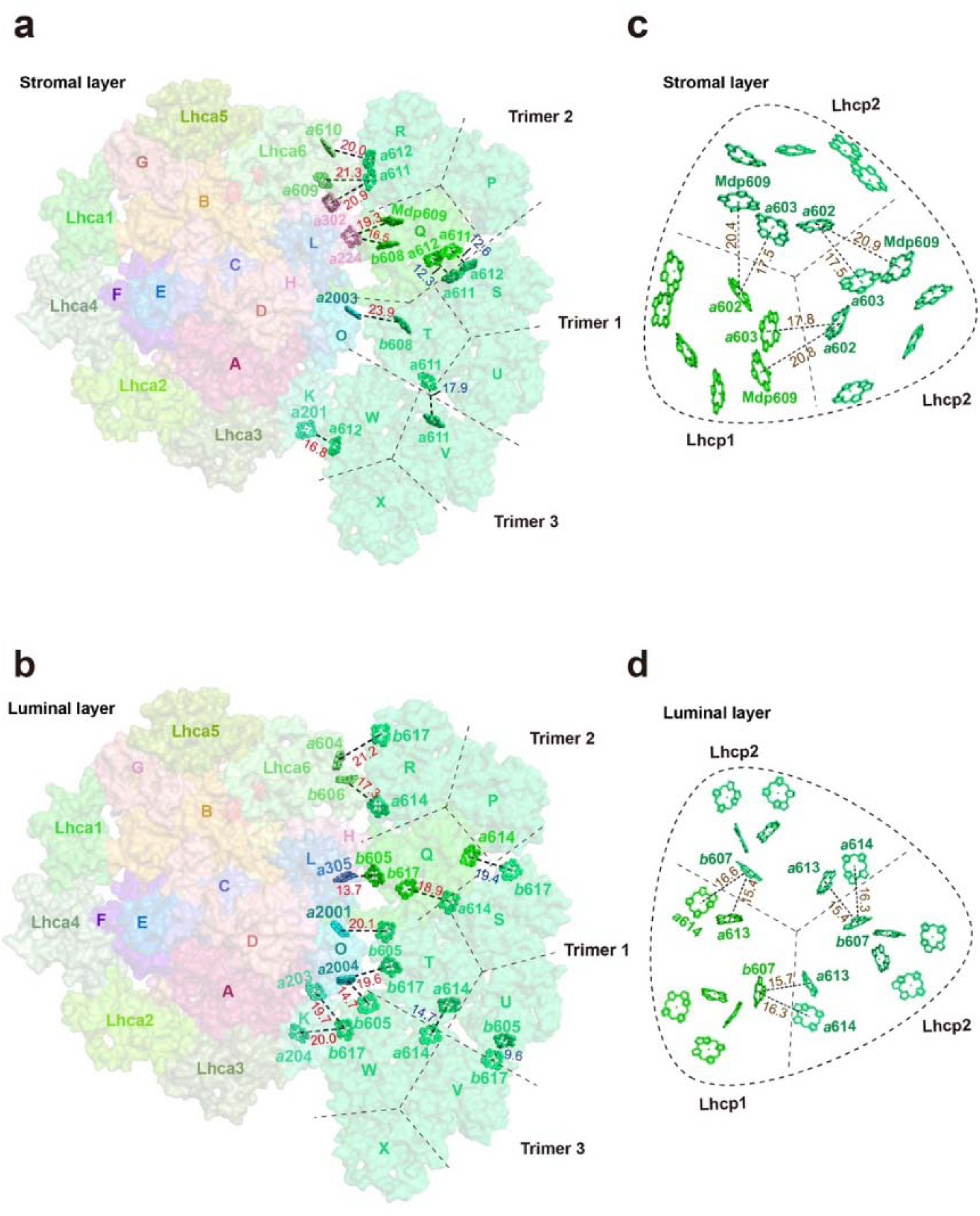
Potential energy transfer pathways between Trimers and PSI. **a** and **b,** The arrangement of interfacial chlorophyll molecules within the stromal (a) and luminal (b) layers. The view is from stromal side and approximately along membrane normal. The chlorophyll molecules at the interfaces between adjacent Trimers and between PSI and Trimers are highlighted as stick models, while the surface presentation of the supercomplex is shown in the background. **c** and **d**, The arrangement of chlorophyll molecules in the Trimer within the stromal (c) and luminal (d) layers. The chlorophyll molecules at the interfaces between adjacent monomers of Trimer are highlighted as stick models. Color codes: light green, Lhcp1; mint green, Lhcp2. The red, blue and brown labels indicate the close connections between two adjacent chlorophyll molecules at the interface between Trimer and PSI/Lhca6 subunits, between two adjacent Trimers and between two adjacent Lhcp monomers within the trimer, respectively. The Mg-Mg distances (Å) are labeled nearby the dotted lines. The dark dashed lines define the estimated interfaces between the adjacent Lhcp monomers, while the dashed round triangles outline the approximate boundary of the trimer . Chains P, Q and R, Trimer 2; Chains S, T and U, Trimer 1; Chains V, W and X, Trimer 3. The phytol tails of chlorophyll molecules are omitted for clarity. **Figure supplement 1.** Structure-based analysis of FRET networks within the PSI-LHC-Lhcp supercomplex.

To supply the excitation energy for PSI, the Chls in Trimer 2 and Trimer 3 make multiple pairwise close connections with the Chls from Lhca6/PsaH and PsaK, respectively (Fig. 10a and b). In contrast, Trimer 1 appears to be poorly connected with PSI as there is only one pair of Chls at their interface in the stromal layer (Fig. 10a), namely Chl *b*608_T/Trimer1_ and Chl *a*2003_PsaO_, and they are separated at a relatively large Mg-Mg distance (23.9 Å). As the distance and orientation factor as well as the involved Chl species determine the efficiency of the inter-complex energy transfer(Croce & van Amerongen, 2020), three Chl pairs, namely Chl *a*612_W/Trimer3_-Chl *a*201_PsaK_, Chl *a*612_R/Trimer2_ -Chl *a*610_Lhca6_, and Chl *a*611_R/Trimer2_ -Chl *a*302_PsaH_, are predicted to mediate the most efficient energy transfer from Trimers to PSI (Table 1), whereas no Chl *a* in Trimer 1 was involved. Although the Chl *a*610_W/Trimer3_-Chl *a*201_PsaK_ or the Chl *a*611_R/Trimer2_-Chl *a*609_Lhca6_ distance is also short (19.2 or 21.3 Å), their energy transfer efficiency would be negligible due to the relatively low orientation factor. Thus, most of the energy collected by Trimer 1 might be transferred to Trimer 2 or Trimer 3 and then pass through Chl *a*611-Chl *a*612_R/Trimer2_ or Chl *a*612_W/Trimer3_ to reach PSI core (Figure 10—supplement 1). Both Trimer 2 and Trimer 3 establish close connections with PSI subunits (PsaH and PsaK) through the Chl *a*611-*a*612 clusters from monomers R and W, respectively. In plant LHCII, the excitation energy within a trimer is mainly populated on the Chl *a*610-*a*611-*a*612 cluster upon final equilibrium(Novoderezhkin et al., 2005). The Chl *a*610-*a*611-*a*612 clusters in monomer R_Trimer2_ and monomer W_Trimer3_ may have a similar function in collecting energy from the adjacent Lhcp monomers and then transfer the energy to the nearby Chls on PsaH and PsaK. Meanwhile, Lhca6 may also serve as an intermediate complex connecting the energy transfer process from Trimer 2 to PSI.

### Physiological role of the *Ot*PSI–LHCI–Lhcp supercomplex

The structure of *Ot*PSI–LHCI–Lhcp supercomplex revealed in this study shows the association of Lhcp trimers with the PSI core and the interfacial pigment pairs, which supports their function as peripheral antennae. Interestingly, the LHCII trimer(s) bound to the same side of the PSI core have been known as conditional PSI antennae in *C. reinhardtii* and higher plants under state 2 conditions (Huang et al., 2021; Pan et al., 2018; Pan et al., 2021). The *O. tauri* cells in the present study were, however, grown in white LL (50 μE m^-2^ sec^-1^), not in a condition known to induce state 2 in *C. reinhardtii* or in higher plants. We compared the amount of A3L band in the cells grown under LL and HL conditions (Fig. 11a). The amount of A3L was more abundant in LL than HL, suggesting that the present PSI–LHCI–Lhcp supercomplex is a LL-acclimated form in this primitive alga, which was also consistent with the previous study(Swingley et al., 2010).

**Fig. 11.**
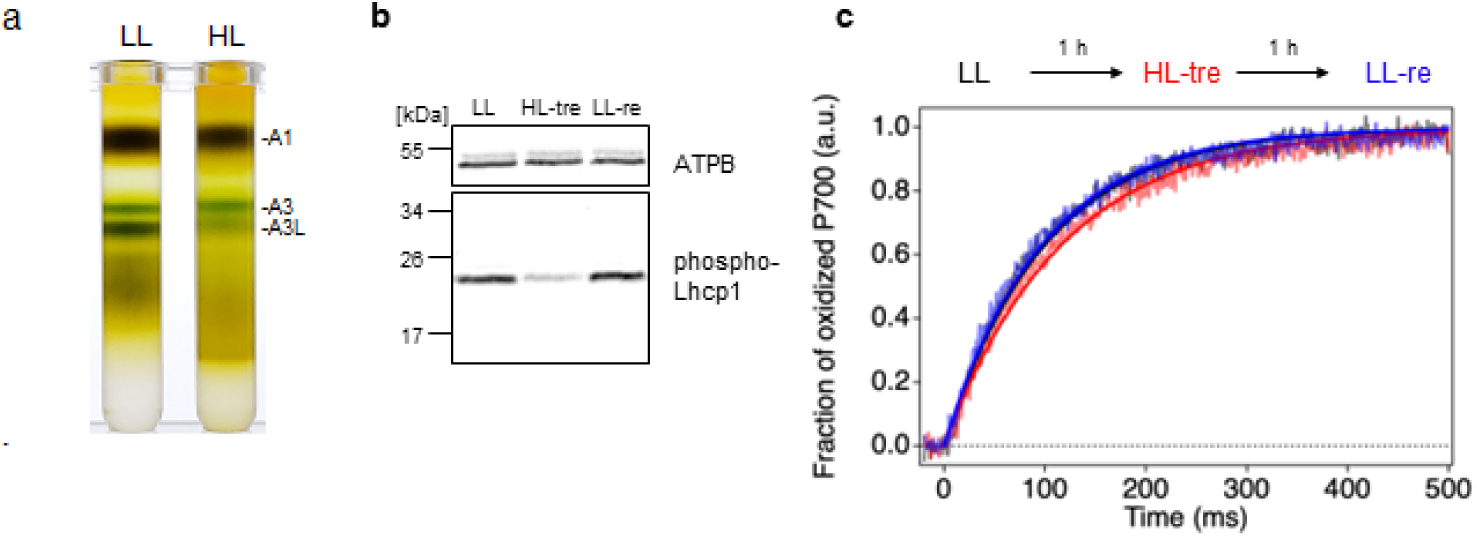
Long- and short-term acclimation of PSI supercomplex in *O. tauri.* **a**, Sucrose density gradient ultracentrifugation of the solubilized thylakoids (0.25 mg Chl) from *O. tauri* cells grown under LL and HL conditions. **b**, Immunoblot analysis of phospho-Lhcp1. Thylakoids (2 µg Chl) were isolated from LL-, HL-treated, and LL-recovered cultures (LL (*black*), HL-tre (*red*) and LL-re (*blue*), respectively). ATPB protein levels are shown as the loading control. Data are representative of two independent experiments. See another set of data in Figure 11—supplement 1, **c**, Light-induced P700 oxidation kinetics measured under 22 µmol photon m^-2^ s^-1^ in the thylakoids sampled from the same cultures in **b**. The fraction of P700 oxidation was derived from Δ(A_820_-A_870_). *Solid* lines and *shaded* lines represent fitting curves by mono-exponential functions and averaged lines of measurements of eight technical replicates, respectively. Data are representative of two independent experiments. See another set of data in Figure 11—supplement 2. **Source data 1.** Quantitative data for ***Figure 11b***. **Source data 2.** Raw data for ***Figure 11c***. **Figure supplement 1.** Immunoblot analysis of phospho-Lhcp1. **Figure supplement 1—source data 1.** Raw data for Figure 11***—figure supplement 1***. **Figure supplement 2.** Light-induced P700 oxidation kinetics measured under 22 µmol photon m^-2^ s^-1^. **Figure supplement 2—source data 1.** Quantitative data for Figure 11***—figure supplement 2*.**

Because the overall architecture of the docking site between the phosphorylated Lhcp1_Trimer 2_ and PsaH/L in *O. tauri* is resembles those reported in the phosphorylated Lhcb2 and PsaH/L in *Z. mays*(Pan et al., 2018) and the phosphorylated LhcbM1 and PsaH/L in *C. reinhardtii*(Pan et al., 2021) including the RRpT motif (Fig. 6d), the mechanism behind this LHC trimer docking event might be similar to that of state transitions(Rochaix, 2014). During state transitions, LHCII trimers are phosphorylated when the plastoquinone pool becomes reduced, and a portion of the phospho-LHCII trimers associate with the pocket formed by PsaH/L on PSI core using the RRpT motif. Since state transitions are commonly observed in both branches of green lineages, namely chlorophytes including *C. reinhardtii* and streptophytes including *Z. mays*, it is possible that a primitive form of state transitions operates in *O. tauri*, whose ancestors sit near the branch point of chlorophytes and streptophytes (Lewis & McCourt, 2004). Therefore, we conducted the following in vivo experiments to test the possibility that the observed acclimation of the PSI supercomplex in the long-term (Fig. 11a) may in fact occur in the short term according to the state transition-like mechanism. The *O. tauri* cells grown under LL conditions (LL-culture) were incubated under HL for 1h (HL-treated culture), and then returned to LL conditions for another 1h incubation (LL-recovered culture). The phosphorylation of Lhcp1 was immunologically detected and the deformation of PSI–LHCI–Lhcp supercomplex was monitored by analyzing the decrease of the PSI antenna size. The Lhcp1 phosphorylation level was greatly reduced when the culture was treated by HL, while it was recovered upon the culture was returned to LL (Fig. 11b, Figure 11—supplement 1), indicating that the Lhcp phosphorylation/dephosphorylation is regulated in the short term depending upon light intensity. It is possible that the mechanism behind this short term event is common to state transitions, but future research in this direction is much expected. Strikingly, the PSI antenna size was in fact reduced corresponding to the Lhcp1 dephosphorylation when the LL-culture was treated by HL, although the reverse reaction, namely restoration of the antenna size, was unclear, which suggests that some unknown factor(s) is possibly involved in this reaction (Figure 11—supplement 2). Together, these results suggest that the antenna size of PSI–LHCI supercomplex in *O. tauri* is subjected to short-term acclimation depending upon light intensity, which is possibly based on a regulatory mechanism similar to state transitions. Such a mechanism may help to increase the light-harvesting capacity of PSI by recruiting the Lhcp trimers under LL condition, and to protect PSI from photodamage by dissociating the Lhcp trimers and reducing the input of excitation energy under HL conditions.

Future work will allow elucidating the relationship between the long-term acclimation that we initially observed (Fig.11a) and the short-term acclimation that suggests a state transition-like event occurring (Fig.11b-c). Several other open questions are to be clarified in future works. For instance, how do LL and HL get the plastoquinone pool reduced and oxidized to regulate the phosphorylation level of Lhcp1, respectively, in this alga? Which kinase and phosphatase in *O. tauri* chloroplast are responsible for phosphorylation and dephosphorylation of Lhcp1, and how are their activities regulated in response to light intensity change? Whether the PSII antenna size varies complementary to that of the PSI? Should we accept this short-term acclimation as state transitions after all?

In conclusion, the cryo-EM structure of PSI supercomplex in *O. tauri* revealed hybrid features of the plant-type and the green algal-type PSI supercomplexes with three Lhcp trimers at the ‘state 2’ position. Our structural, biochemical and spectroscopic analysis results collectively suggest that *O. tauri*, the early branching green alga, may develop a light intensity-induced state transition-like mechanism to regulate the light-harvesting process.

## MATERIALS AND METHODS

### Strain and growth conditions

*O. tauri* strain OTH95 (Derelle et al., 2006) was obtained from Roscoff Culture Collection. Cells were grown in artificial seawater made with Sigma sea salt (Sigma cat#59883) supplemented with Daigo’s IMK medium (Fujifilm-Wako, Japan) at 25 °C under white fluorescence bulb (Panasonic FL40SS W/37R) at the indicated light intensities with constant air-bubbling and gentle stirring. For the short-term photoacclimation experiments, cells grown at 50 µmol photon m^-2^s^-1^ (LL-culture) were shifted to 500 µmol photon m^-2^s^-1^ for 1 h (HL-treated culture), then returned to 50 µmol photon m^-2^s^-1^ and incubated for 1 h (LL-recovered culture).

### Preparations of pigment-protein complexes

Thylakoid membranes were prepared as described previously(Swingley et al., 2010), with a modified cell lysis protocol by using two rounds of nebulization by BioNeb (Glas-Col, Terre Haute, IN) at 8 kg/cm^2^ force (of N_2_). The obtained membranes were solubilized with dodecyl-α-D-maltoside (Anatrace, Maumee, OH), which was subsequently replaced by amphipol A8-35 (Anatrace), and the pigment-protein complexes were fractionated by the sucrose density gradient ultracentrifugation as described previously(Watanabe et al., 2019).

### SDS-PAGE and Immunoblotting

SDS-PAGE and Immunoblotting were performed as described previously(Iwai et al., 2008). Anti-phospho-Lhcb2 (AS13 2705) and anti-ATPB (AS05 085) antibodies were obtained from Agrisera (Vännäs, Sweden).

### Mass spectrometry

The sample was applied for trypsin digestion(Shevchenko et al., 2006) and analyzed by a UPLC system (EASY-nLC 1000, Thermo Fisher Scientific) coupled to an Orbitrap Elite mass spectrometer (Thermo Fisher Scientific). The analyzed data were submitted for protein database search using Proteome Discoverer software (Thermo Fisher Scientific) and Mascot version 2.5.1 (Matrix Science). The protein database was generated from the polypeptide sequences of *O. tauri* OTH95 deposited in NCBI(Derelle et al., 2006).

### Pigment analysis

Pigments were analyzed on a UPLC system as described previously(Tokutsu & Minagawa, 2013) with the following modifications. Pigments were extracted from the samples with 80 % aceton. Separation was carried out on a Cadenza CD-C18 UP, 2× 150 mm, 3-µm column (Imtakt, Kyoto, Japan). Gradient elution was established with three-solvent system, acetonitrile/isopropanol/water. The gradient was shifted from 65:15:20 (vol/vol) to 80:15:5 (vol/vol) at 6.5 min and to 50:50:0 (vol/vol) at 10 min before returning to 65:15:20 (vol/vol) at 15 min. The column temperature was 45 °C. The system was calibrated with commercial standards as long as they were available (DHI, Hoersholm, Denmark). Concentrations of Dlt and Mdp were estimated based on the response factor of lutein and Chl *c*_2_, respectively, and uriolide and micromonal were estimated based on the response factor of Prx as previously described(Latasa et al., 2004). We used extinction coefficients from (Roy et al., 2011).

### UV-Vis absorption spectroscopy

Absorption spectra were obtained at 0.5-nm intervals in the range 400 to 800 nm by using a UV-Vis spectrometer V-650 (JASCO Corp, Tokyo, Japan) with an integrating sphere ISV-722 (JASCO) at room temperature. Prior to the measurements, concentration of the samples was adjusted to 3 µg Chl ml^-1^.

### Fluorescence spectroscopy

Time-resolved fluorescence spectra were measured at 77 K as described previously(Yokono et al., 2015), by using a photoluminescence spectrometer FLS1000-ss-stm (Edinburgh Instruments, Livingston, UK). Prior to the measurements, concentration of the samples was adjusted to 4 µg Chl ml^-1^. Excitation wavelength was 405 nm. The instrumental function was measured with each sample at 405 nm with 2 nm bandwidth. Typical FWHM of the instrumental function was 50 ps, and maximum time-resolution after deconvolution analysis was 5 ps. The excitation laser intensity was less than 8 µW with the repetition rate at 2 MHz, which did not interfere with measurements up to 100 ns. Time intervals is 2.44 ps/channel for up to 10 ns and 97.7 ps/channel for up to 100 ns. The optical slit was set to 10 nm and the optical filter (LOPF-25C-593) was used to cut the excitation pulse from the fluorescence signal. The signal-to-noise ratio was over 30,000:1. The fluorescence decay-associated spectra and the steady-state fluorescence spectra were constructed as described previously (Yokono et al., 2015; Yokono et al., 2019). Briefly, the fluorescence decays were fitted using convolution and simulation method using Mathematica 12 (Wolfram Research, Champaign, IL, USA). Instrumental function and free exponential components were used to simulate fitting curves, and the determined exponential components were used to construct deconvoluted decay curves. The deconvoluted decay curves were imported to Igor 8 (WaveMetrics, Lake Oswego, OR, USA) to perform global analyses. Following a global analysis of the fluorescence kinetics, FDAS were constructed. For the PSI–LHCI and the PSI–LHCI–Lhcp supercomplexes, steady-state fluorescence spectra were re-constructed from 1st and 2nd lifetime components of FDAS by integration on time axis.

### PSI antenna size measurements

Thylakoids were prepared as follows. Twenty ml of cell culture was initially shock frozen with liquid N_2_ under the culturing light either at 50 µmol photon m^-2^s^-1^ (LL-culture and LL-recovery-culture) or 500 µmol photon m^-2^s^-1^ (HL-culture). The frozen cells were disrupted by adding 30 ml of disrupting buffer (25 mM HEPES (pH 7.5), 10 mM sodium fluoride, and proteinase inhibitor Pefabloc®SC at 0.25 mg ml^-1^ (Roche, Basel, Switzerland) before centrifugation at 22,000 g for 2 min at 4 °C. The pellets were resuspended in measuring buffer (25 mM HEPES (pH 7.5), 330 mM sucrose, 1.5 mM NaCl, 5 mM MgCl_2_, 10 mM sodium fluoride) at 60 μg Chl ml^-1^. A3 and A3L fractions were resuspended in measuring buffer (25 mM HEPES, pH 7.5) at 20 µg Chl ml^-1^. These samples were incubated in the presence of 100 µM 3-(3,4-dichlorophenyl)-1,1-dimethylurea (DCMU), 5 mM sodium ascorbate and 400 μM methyl-viologen for 5 min to create a donor-limited situation (Melis, 1982) and P700 oxidation kinetics were analyzed by monitoring the difference between two transmission pulse signals at 820 and 870 nm using a Dual/KLAS-NIR spectrophotometer (Heinz Walz GmbH, Effeltrich, Germany) as previously reported (Klughammer & Schreiber, 2016). Dark-light induced kinetics were obtained with red actinic light (16, 28, and 48 µmol photon m^-2^s^-1^, 635 nm). Traces were normalized between the minimum (before actinic light illumination) and the maximum values and fitted with a mono-exponential function to determine the P700 oxidation rate (τ^-1^).

### Negative staining single particle analysis

Negative staining single particle analysis was performed as described previously(Watanabe et al., 2019) with following modifications. Isolated protein samples were diluted to 2 µg Chl ml^-1^. Electron micrographs were obtained at 100,000× magnification and recorded at a pixel size of 5.0 Å. In total, 50 micrographs for the A3, 200 micrographs for A3L were collected. Two dimensional classification was performed into 50 classes. Small PSI (PSI–LHCI) and large PSI (PSI–LHCI–Lhcp) supercomplexes were assigned based on the shape of the averaged particles.

### FRET calculation

FRET rate constants were computationally calculated based on the simplified FRET principle with an approximation that all Chl *a* and *b* had the identical excited state energy levels so the spectral overlap integral for each combination was constant as previously described(Mazor et al., 2017; Sheng et al., 2019).

### Cryo-EM data collection, processing, classification and reconstruction

A total of four datasets were collected on a 300 kv Titan Krios microscope (FEI) equipped with K2 camera (Gatan) by using four different grids prepared under similar conditions. Movies were captured with a defocus value at the range of -1.8 to -2.2 μm. The physical pixel size and total dose are 1.04 Å and 60 e^-^/Å^2^, respectively. After removing images with poor quality or evident drift, a total of 19,680 (2,697, 2,992, 6,870 and 7,121 for each dataset) images were selected and used for further processing. The beam-induced motion in each movie with 32 frames were aligned and corrected using MotionCor2 (Zheng et al., 2017). The corrected images after alignment were used for estimation of the contrast transfer function(CTF) parameters by using CTFFIND4.1(Rohou & Grigorieff, 2015). The initial image-processing steps, including manual particle picking, reference-based particle autopicking, reference-free two-dimensional(2D) classification and three-dimensional(3D) classification, were performed in cryoSPARC(Punjani et al., 2017). The image subsets were divided into about 10 groups by reference-free 2D classification, which were further applied as templates for reference-based particle autopicking. Subsequently, the total of 5,288,217 raw particles were picked and further classified through the 2D classification procedure. As a result, 842,823 particles with good contrast were output from cryoSPARC and imported into Relion 3.1(Scheres, 2012) for further processing step. The particles were subjected to 3D classification without providing any references. The selected classes (2,452, 6,648, 32,751 and 38,722 particles for each of the four datasets) of PSI–LHCI–Lhcp supercomplex from 3D classification were combined and used for further processing through the 3D auto-refinement with *C*1 symmetry, CTF refinement and Bayesian particles polishing. The final cryo-EM density map resolution of *Ot*PSI–LHCI–Lhcp supercomplex was estimated by using the gold standard Fourier shell correlation (FSC) of two half maps with a cutoff at 0.143. To identify each individual subunits of the three Trimers, the local refinement procedure was carried out to further improve the map quality around each of the three Trimers. The local masks around each individual Trimers were applied during the refinement and the final resolutions of Trimers 1, 2 and 3 were 3.3 Å, 3.4 Å, 2.9 Å, respectively.

### Model building and refinement

To build the model of *Ot*PSI–LHCI–Lhcp supercomplex, the structure of *Zm*PSI–LHCI (PDB code: 5ZJI) was fitted into the PSI–LHCI region of the 2.94-Å cryo-EM map by using UCSF chimera and manually adjusted through rigid body fit zone in COOT to achieve improved matching of the model with density. Subsequently, the amino acid residues in the model of each chain were corrected and registered by referring to the sequences of corresponding proteins from *Ostreococcus tauri*. The PsaM subunit was added manually by *de novo* model building with the amino acid sequence as a reference. The Lhca5 and Lhca6 model was built by referring to the corresponding ones from the *Cr*PSI–LHCI–LHCII structure(PDB code: 7D0J) and each amino acid residues in the model were checked and corrected by referring to local density feature and the *Ot*Lhca5 and *Ot*Lhca6 sequences. The density of Lhcp1 exhibits an elongated N-terminal region similar to the corresponding part in *Cr*LhcbM1, and shows a well-defined side chain densities in the local part found in the surface binding pocket of PSI. The model of Lhcp1 was built by using *Cr*LhcbM1 (PDB code: 7D0J) as an initial model, sequence registration and correction of the model was performed by referring to the map and *Ot*Lhcp1 sequence simultaneously. For the model of Lhcp2, an initial model was adapted from an LHCII monomer, which was corrected and refined according to the cryo-EM map and *Ot*Lhcp2 sequence. For pigment molecules, two different carotenoids (Prx and Dlt), and one Chl (Mdp) found in prasinophyte were identified and assigned in each Lhcp1/2 monomer by referring to their characteristic density features. As a result, each Lhcp1/2 monomer contains 4 Dlt, 2 Prx and 1 Mdp in addition to 1 Nex, 8 Chl *a* and 5 Chl *b*. Among the Chls, one (Chl *b*617) is located at the peripheral region of each Lhcp1/2. These pigment molecules show well-defined density features useful for their identification and model building. As the unambiguous identification of Chl *a* from Chl *b* requires higher resolution at around or beyond 2.8 Å, the current models of Chl *a* and Chl *b* were tentatively assigned by considering the local features and binding environments around C7 group.

After model building, real-space refinement of the *Ot*PSI–LHCI–Lhcp supercomplex structure against the 2.94-Å overall map was performed by using Phenix 1.19.2 (Afonine et al., 2018). The models of Trimers 1 and 3 were refined by using the 3.3-Å and 2.9-Å local maps, respectively. The two Lhcp2 (Chain P and Chain R) of Trimer 2 were firstly refined against the 3.4-Å local map, combined with the Lhcp1 model and then further refined against the overall map. The geometric restraints, between magnesium ions of chorophyll molecules or irons of Fe_4_S_4_ clusters and their coordination ligands, were applied during the refinement process. After real-space refinement, manual adjustment and correction was carried out iteratively in COOT. The geometries of the structural model were assessed using Phenix and the detailed information were summarized in Extended Data Table 2.

## Data availability

The atomic coordinates of the *Ot*PSI–LHCI–Lhcp supercomplex has been deposited in the Protein Data Bank with accession code 7YCA. The cryo-EM map of the supercomplex has been deposited in the Electron Microscopy Data Bank with accession code EMD-33737.

## Author contributions

Asako Ishii, Validation, Investigation (biochemical preparation and characterization of the PSI–LHCI–Lhcp supercomplex), Data curation, Visualization; Jianyu Shan, Software: programming, Investigation (building, refinement, and analysis of the structural models), Data curation, Writing—original draft, Visualization; Xin Sheng, Software: programming, Investigation (collection and refinement of cryo-EM data, and model building), Data curation, Visualization; Eunchul Kim, Methodology, Software: programming, Validation, Formal analysis, Investigation (absorption spectroscopy including P700 oxidation kinetics, single particle analysis of the negative staining EM images, and computational analysis on energy transfer), Data curation, Writing—original draft, Visualization; Akimasa Watanabe, Conceptualization, Investigation (biochemical preparation); Makio Yokono, Methodology, Investigation (thylakoid preparation for short-term photoacclimation experiments and fluorescence spectroscopy including FDAS analysis), Writing—original draft; Chiyo Noda, Investigation (pigment analysis and negative staining-EM); Chihong Song: Investigation (prepation of cryo-EM grids); Kazuyoshi Murata, Methodology, Supervision; Zhenfeng Liu, Conceptualization, Methodology, Resources, Data curation, Writing—original draft, Writing—review & editing, Visualization, Supervision, Project administration, Funding acquisition; Jun Minagawa, Conceptualization, Methodology, Formal analysis, Resources, Writing—original draft, Writing—review & editing, Visualization, Supervision, Project administration, Funding acquisition.

## Acknowledgements

We thank Bo-Ling Zhu, Xiao-Jun Huang, Xu-Jing Li and other staff members for their support in cryo-EM data collection at the Center for Biological Imaging (CBI), Core Facilities for Protein Science at the Institute of Biophysics, Chinese Academy of Sciences. We are grateful to Mr. Masato Kubota for his technical assistance for biochemical characterizations and insightful discussion. We also thank Mr. Hiroki Mizoguchi and Yoshihiro Kawada for their involvement in the early phase of this research, Ms. Xiao-Bo Liang for her assistance in sample shipment and handling and Dr. Mei Li for discussion. We are grateful to Mrs. Tomoko Mori and Ms. Yumiko Makino (Trans-Omics Facility, NIBB Trans-Scale Biology Center) for providing technical assistance with LC-MS/MS analysis. The project is funded by Japan Society for the Promotion of Science (JSPS) KAKENHI (21H04778 and 21H05040 to J.M.), the Chinese Academy of Sciences (the Strategic Priority Research Program XDB37020101 and Young Scientists in Basic Research YSBR-015 to Z. L.), the National Natural Science Foundation of China (31925024 to Z.L.) and the National Key R&D Programme of China (2017YFA0503702 to Z.L.). This work was also supported by Model Plant Research Facility, NIBB and the Cooperative Study Program of National Institute for Physiological Sciences (NIPS).

---

**Figure 1—figure supplement 1.**
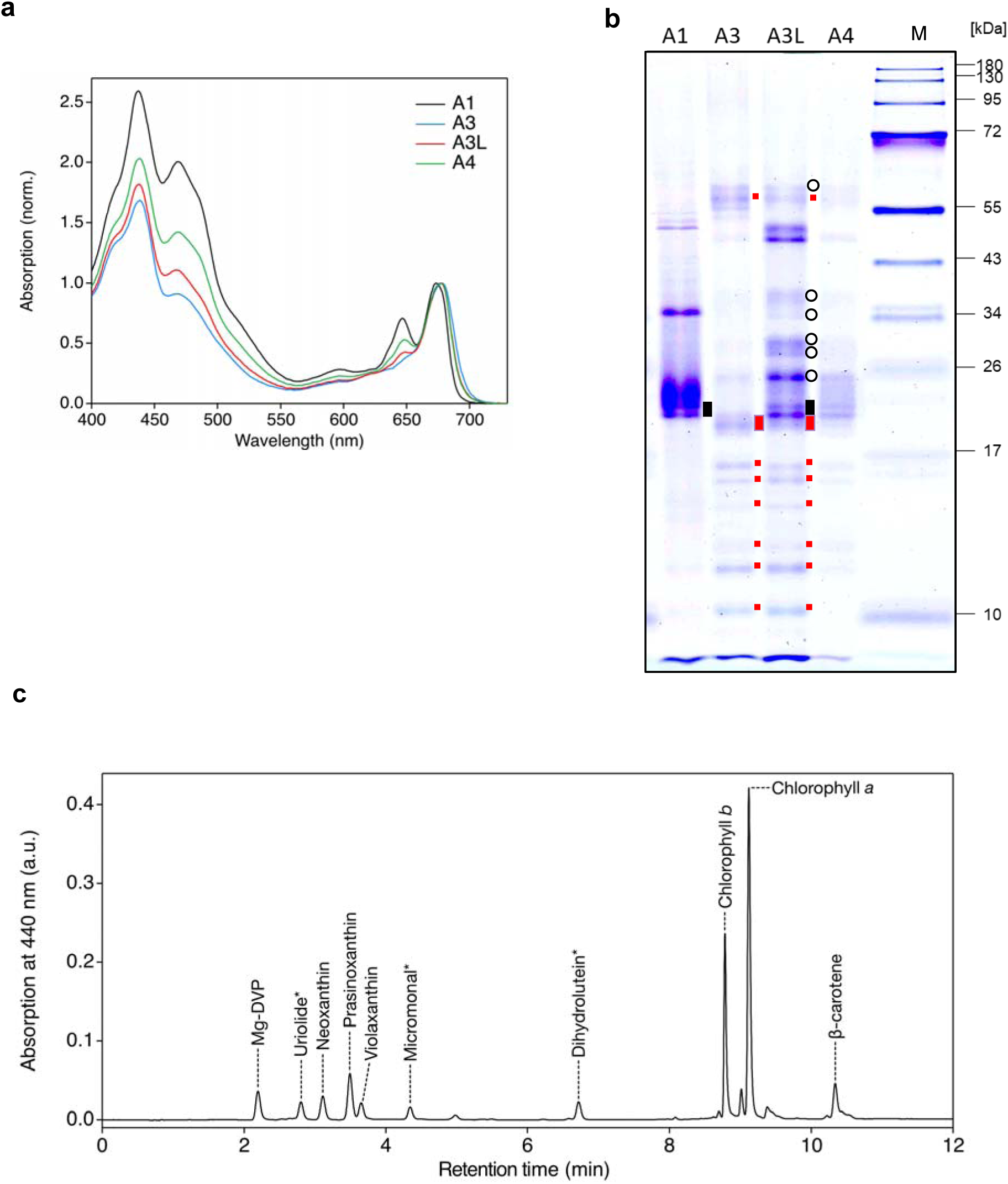
Fractionation and characterization of the supercomplex samples from the *O. tauri* cells grown in the low light (50 µmol photon m^-2^s^-1^). a, UV-VIS absorption spectra. Each fraction (3 μg Chl/mL) was measured at room temperature at least three times and the representative spectra are shown. **b**, SDS-PAGE analysis of each fraction (1 μg Chl/mL) stained with Coomassie Brilliant Blue R-250. M, molecular weight marker. red dots, PSI subunits; black open circles, PSII subunits; black rectangular, Lhcp; red rectangular LHCI. The experiment was repeated three times independently with similar results. **c.** Pigment analysis of the A3L fraction by UPLC. The chromatogram was recorded at 440 nm. Each peak was identified by using a corresponding standard or based on absorption spectrum measured by a photo-diode array equipped in the UPLC system. Asterisks indicate the pigments identified by absorption spectra.

**Figure 1—figure supplement 2.**
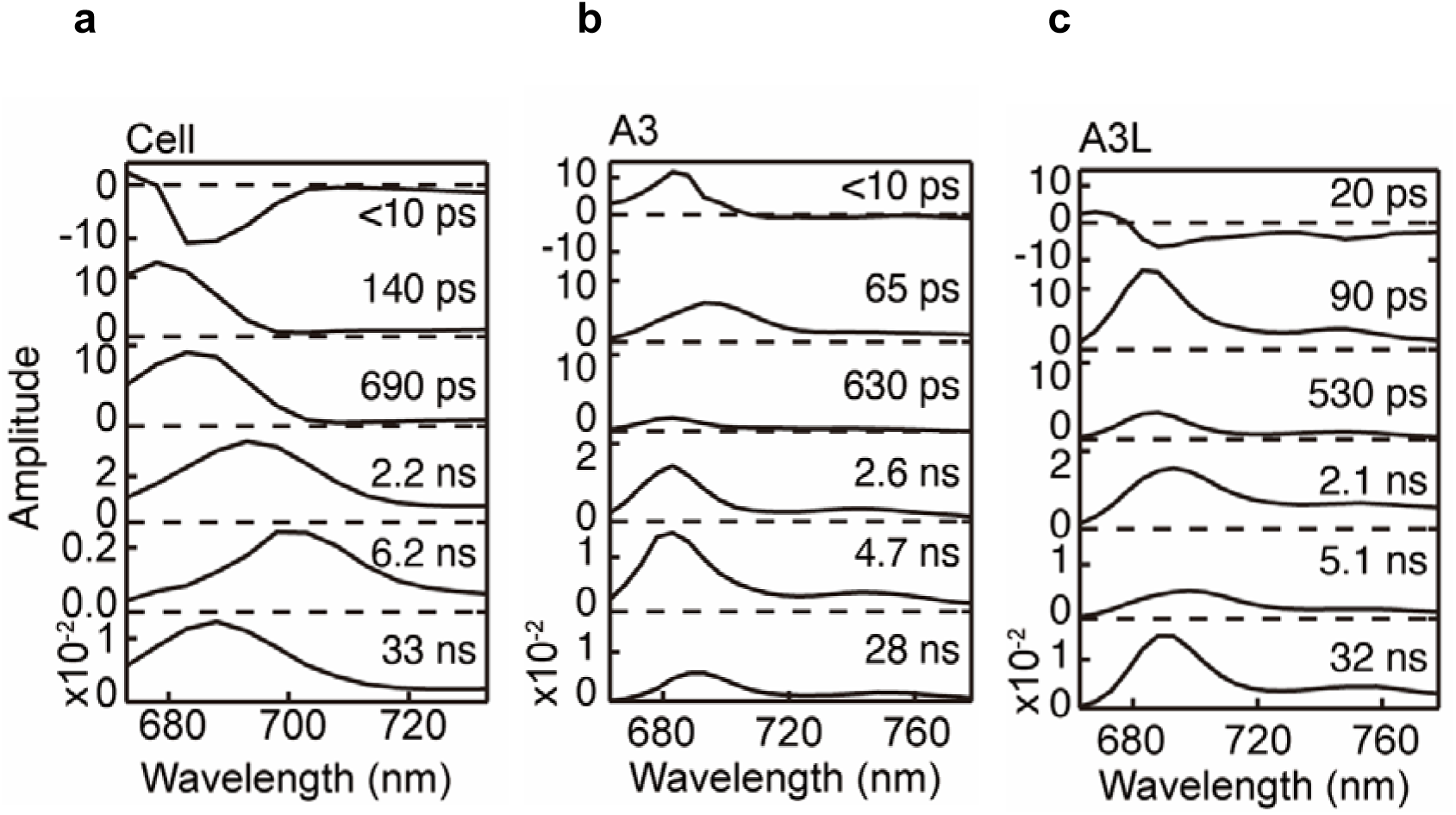
Fluorescence decay-associated spectra (FDAS) at 77K (ex., 405 nm, 4 µg Chl ml^-1^). **a**, A1 fraction. **b**, A3 fraction. **c**, A3L fraction. Fluorescence decay kinetics were fitted to a sum of exponentials with the time constants linked in a global analysis. The normalized amplitudes of FDAS are plotted against wavelength for each time constants (lifetime component).

**Figure 1—figure supplement 3.**
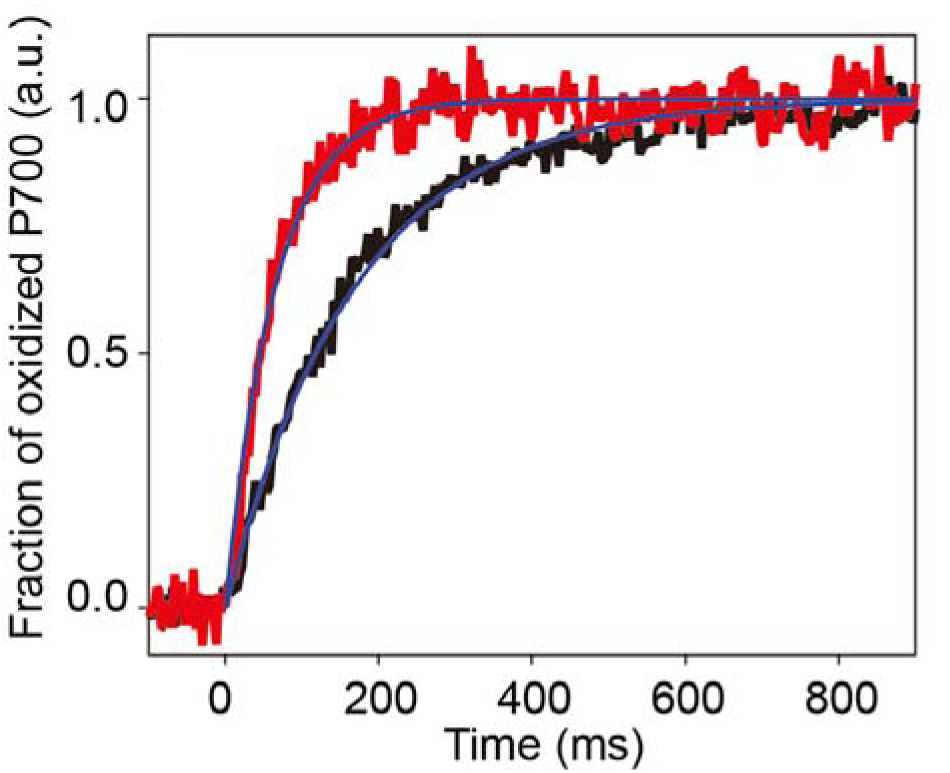
PSI light-harvesting capabilities in the A3 and A3L fractions. All conditions were the same as in Fig.1c.

**Figure 1—figure supplement 4.**
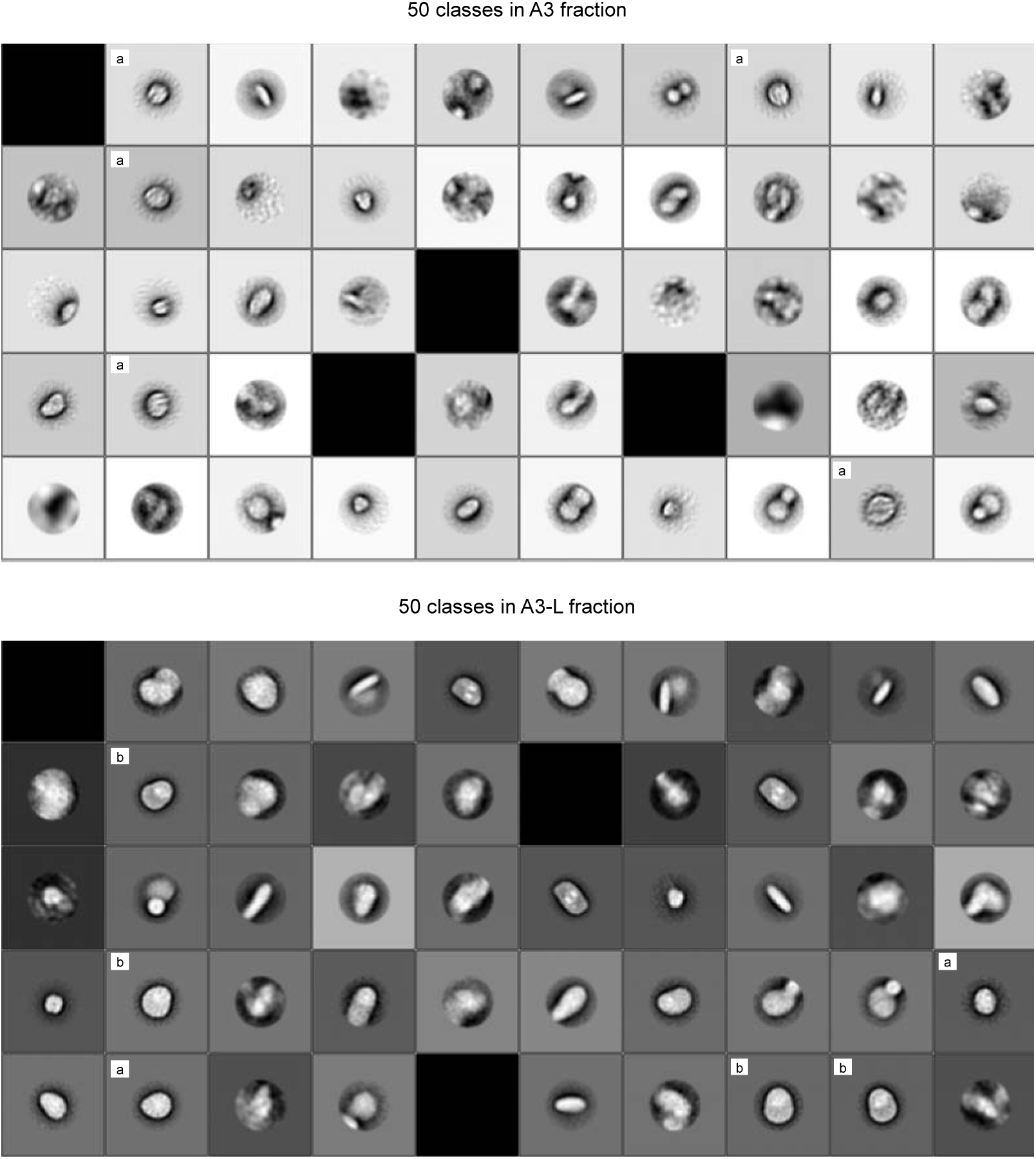
Structural analysis of A3 and A3L fractions by negative staining EM. Th e particles in the A3 and A3L were negatively stained with uranyl acetate and the projection images of EM single particles were 2D classified (Table S3). While most of the PSI in A3 was smaller supercomplex, 70% of the PSI in A3L were larger supercomplex. Two-dimensional classification results of the particles from the A3 and A3L fractions are shown. 9068 particles from the A3 fraction and 10331 particles from A3L were classified into 50 classes. **a** and **b** labels represent classes categorized into small PSI, large PSI, respectively.

**Figure 2—figure supplement 1.**
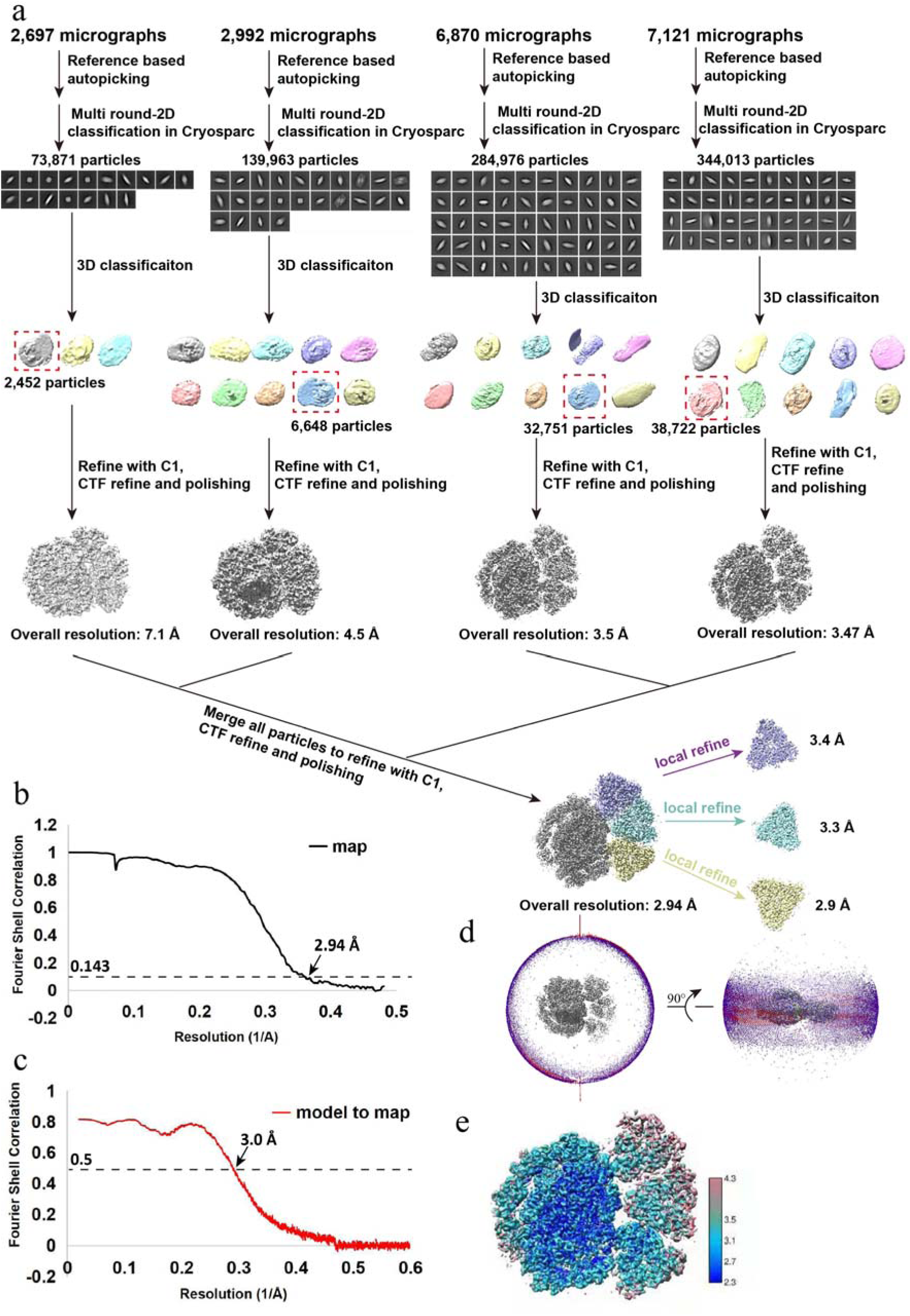
Cryo-EM data collection, processing, refinement and validation statistics of *Ot*PSI-LHCI-Lhcp structures. **a,** Data collection, processing and refinement scheme. **b,** The gold standard Fourier shell correlation (FSC) curves of the final map with criterion of 0.143. **c,** The FSC curves of the refined models versus the cryo-EM map with criterion of 0.5. **d,** Angular distribution of the particles used for the final 3D reconstruction. **e**, Estimation of local resolution of the final cryo-EM map.

**Figure 2—figure supplement 2.**
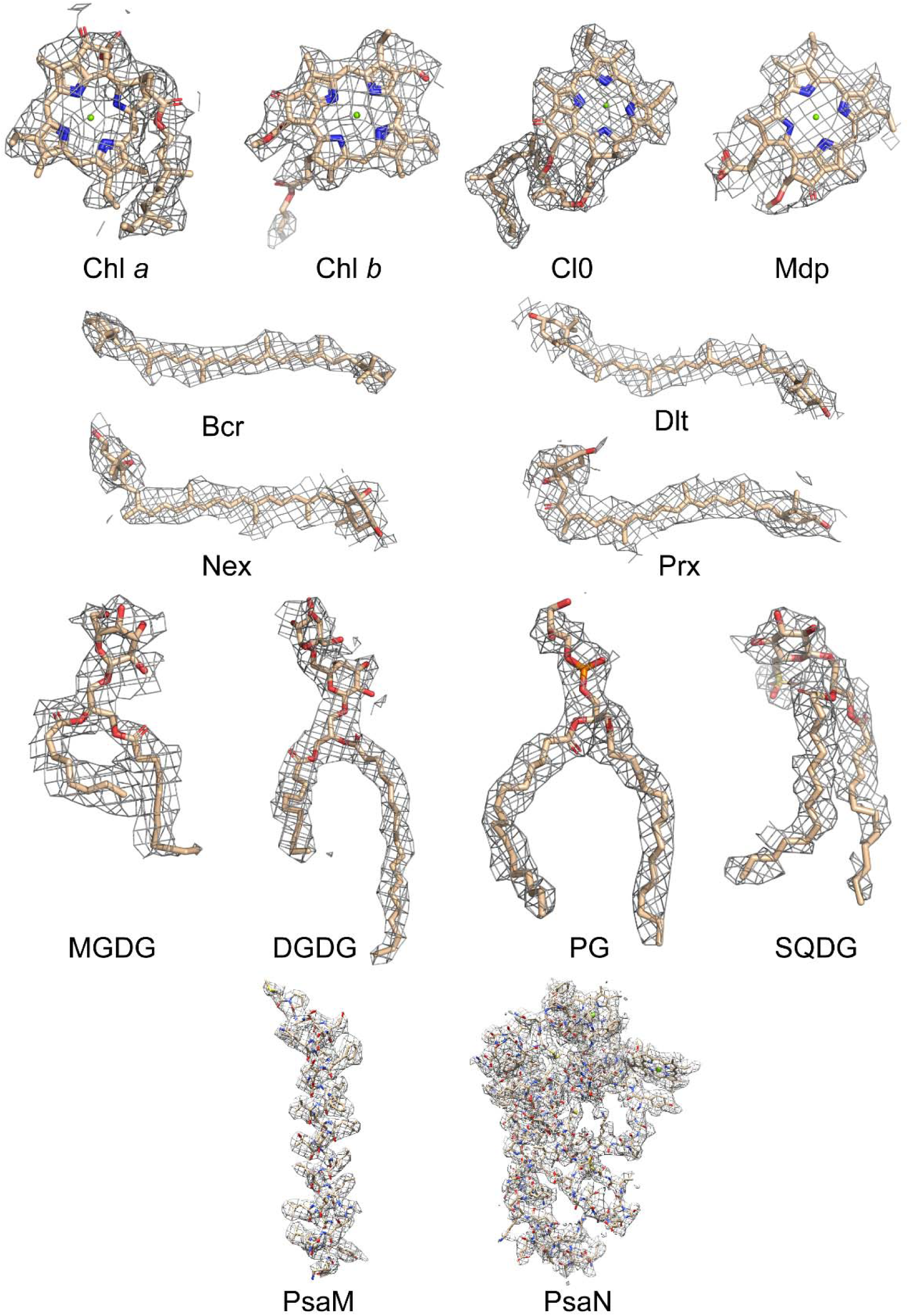
Cryo-EM densities of various cofactors and protein subunits found in the PSI-LHCI-Lhcp supercomplex of *O. tauri.* CLA, chlorophyll. *a*; CHL, chlorophyll *b*; CL0, a chlorophyll *a* isomer of the special pairs of Chl *a* in the reaction center; DVP, Mg-2,4,-divinyl-phaeoporphyrin a5 monomethyl ester; BCR, beta-carotene; DLT, dihydrolutein; NEX, neoxanthin; PRX, prasinoxanthin; MGDG, monogalactosyl-diacylglycerol; DGDG, digalactosyl-diacylglycerol; PG, 1,2-dipalmitoyl-phosphatidylglycerol; SQDG, sulfoquinovosyl diacylglycerol.

**Figure 2—figure supplement 3.**
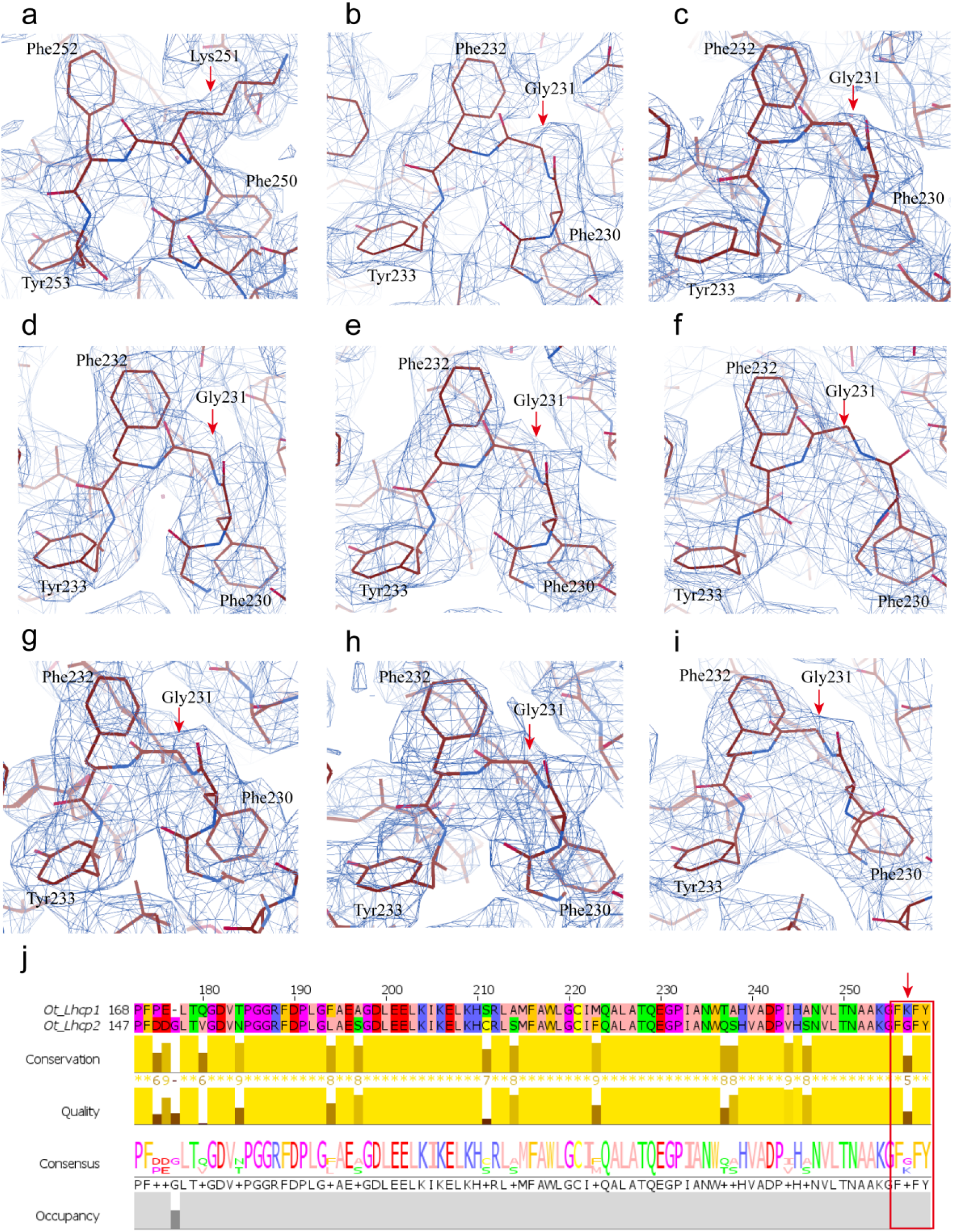
The detailed local cryo-EM map features for identification of *Ot*Lhcp1 a nd *Ot*Lhcp2. Different map features of a C-terminal characteristic motif matches the corresponding FkFY and FgFY models of *Ot*Lhcp1 and *Ot*Lhcp2 respectively. **a-c**, Chain Q, P, R of Trimer1 respectively. **d-f**, Chain S, T, U of Trimer2 respectively. **g-i**, Chain V, W, X of Trimer 3 respectively. The unique residues used for identification are indicated by red arrows. **j**, Sequence alignment analysis of *Ot*Lhcp1 and *Ot*Lhcp2. Note that the amino a cid residue at position 251 of OtLhcp1 is Lys, whereas the corresponding residue at position 231 of OtLhcp2 is Gly.

**Figure 7—figure supplement 1.**
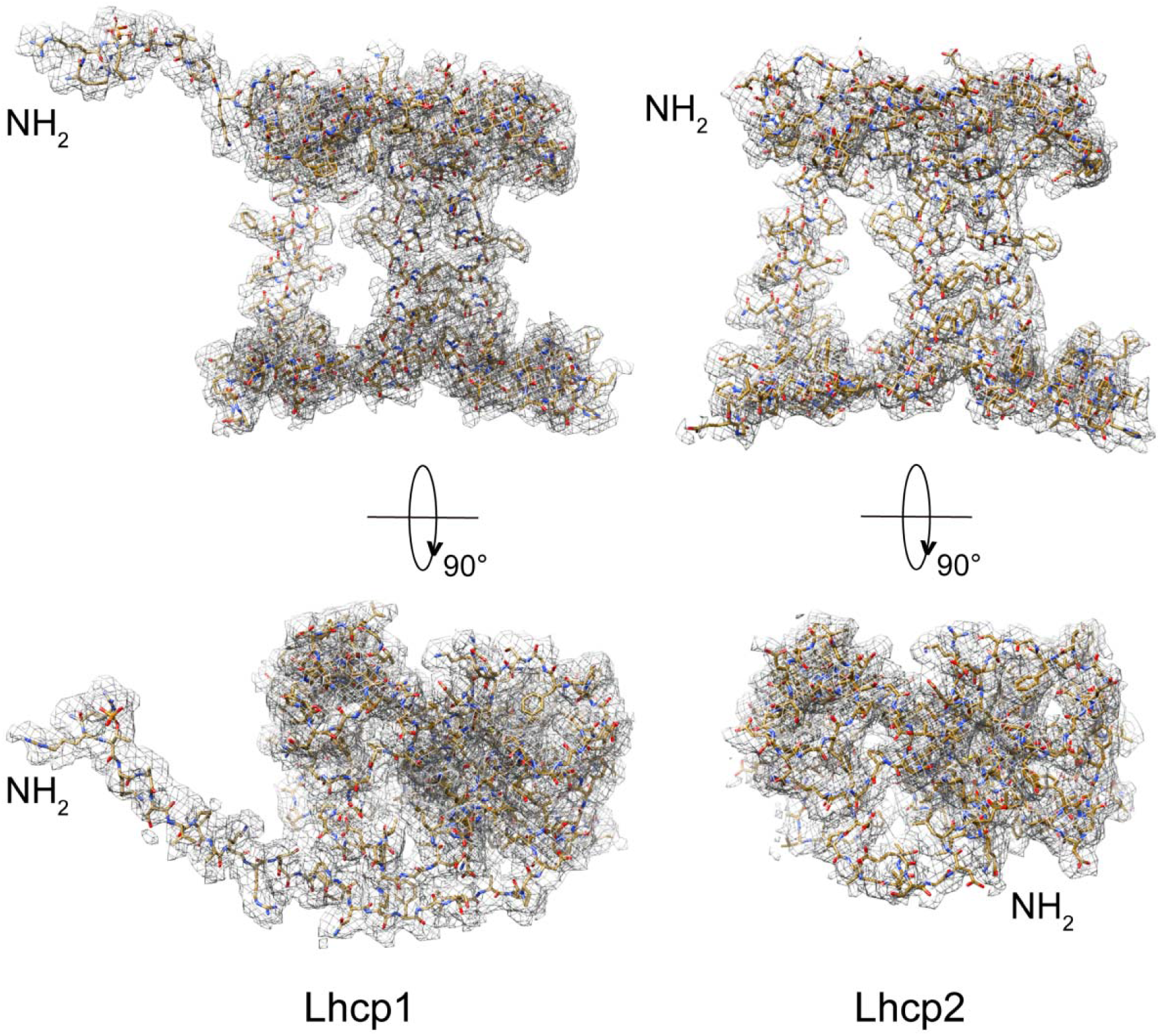
Cryo-EM densities of *Ot*Lhcp1 and *Ot*Lhcp2. The overall structure of *Ot* Lhcp1 highly resembles that of *Ot*Lhcp2, except that the former one has an elongated N-terminal region with a phosphorylated Thr residues. The cofactors are omitted for clarity.

**Figure 7—figure supplement 2.**
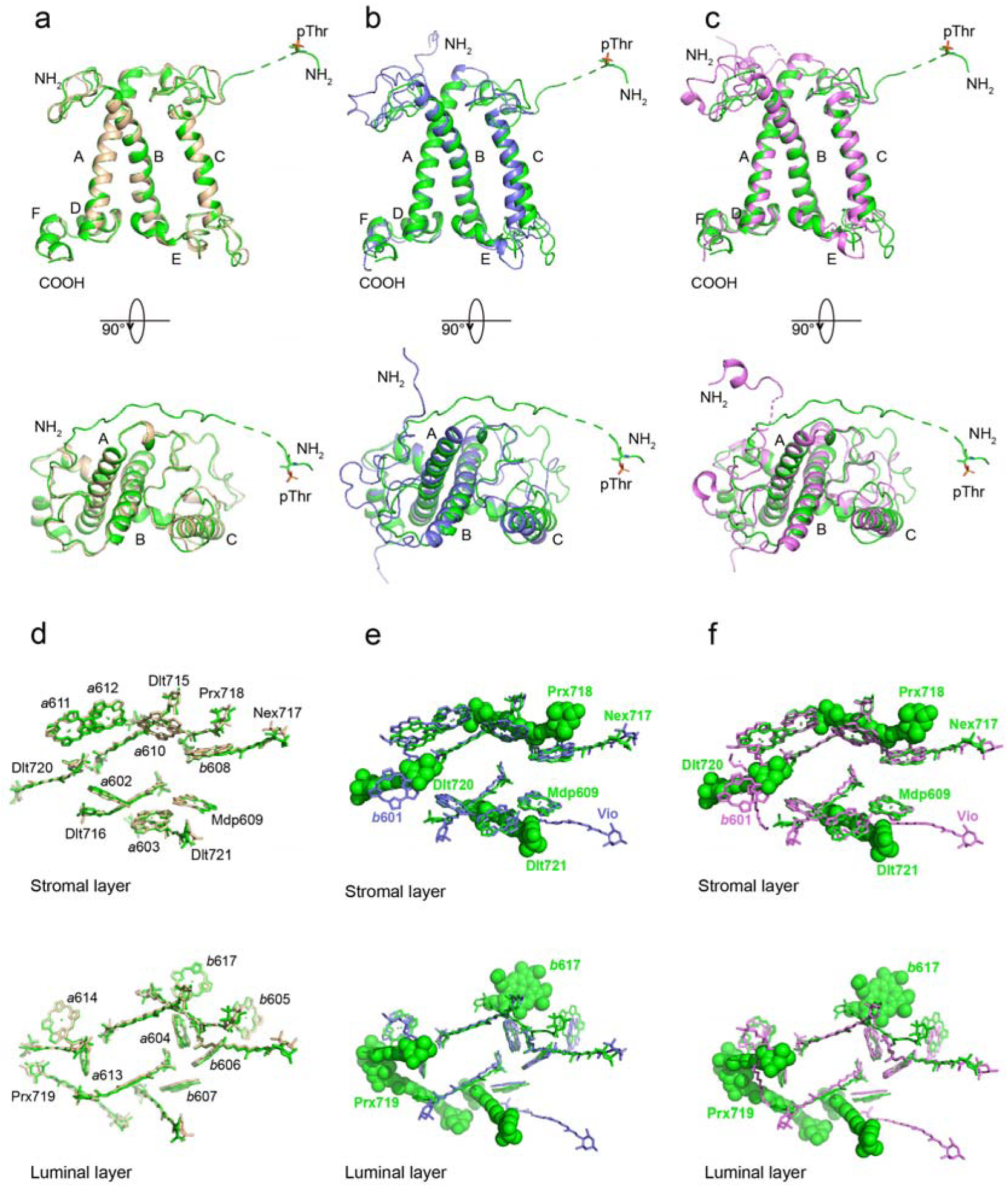
Superposition of the *Ot*Lhcp1 structure with those of *Ot*Lhcp2, *Zm*Lhc b1 and *Cr*LhcbM1. **a**-**c**, The apoprotein of *Ot*Lhcp1 superposed with those of *Ot*Lhcp2 (a), *Zm*Lhcb1 (b) an d *Cr*LhcbM1 (c). Color codes: green, *Ot*Lhcp1; golden, *Ot*Lhcp2; light blue, *Zm*Lhcb1; violet, *Cr*LhcbM1. T he N-terminal phosphorylated Thr residues are highlighted as stick models. A, B and C indicate the three transmembrane helices in Lhcp/Lhcb apoproteins. Upper row, side view along membrane plane; lower row, top view from stromal side along membrane normal. The dash lines indicate the flexible linker regions not observed in cryo-EM maps. **d-f**, The distribution of pigment molecules in *Ot*Lhcp1 in comparison with those in *Ot*Lh cp2 (d), *Zm*Lhcb1 (e) and *Cr*LhcbM1 (f). In the upper row, only chlorophyll molecules within the layer close to stromal surface are shown, while the lower row shows the chlorophyll molecules within the layer close to t he luminal surface. In **e** and **f**, The chlorophyll and carotenoid molecules only present in *Ot*Lhcp1 but not in *Z m*Lhcb1 or *Cr*LhcbM1 are highlighted as sphere models. Those present in *Zm*Lhcb1 and *Cr*LhcbM1 but absent in *Ot*Lhcp1 are indicated by the light blue or violet labels (LHG, *b*601 and Vio).

**Figure 8—figure supplement 1.**
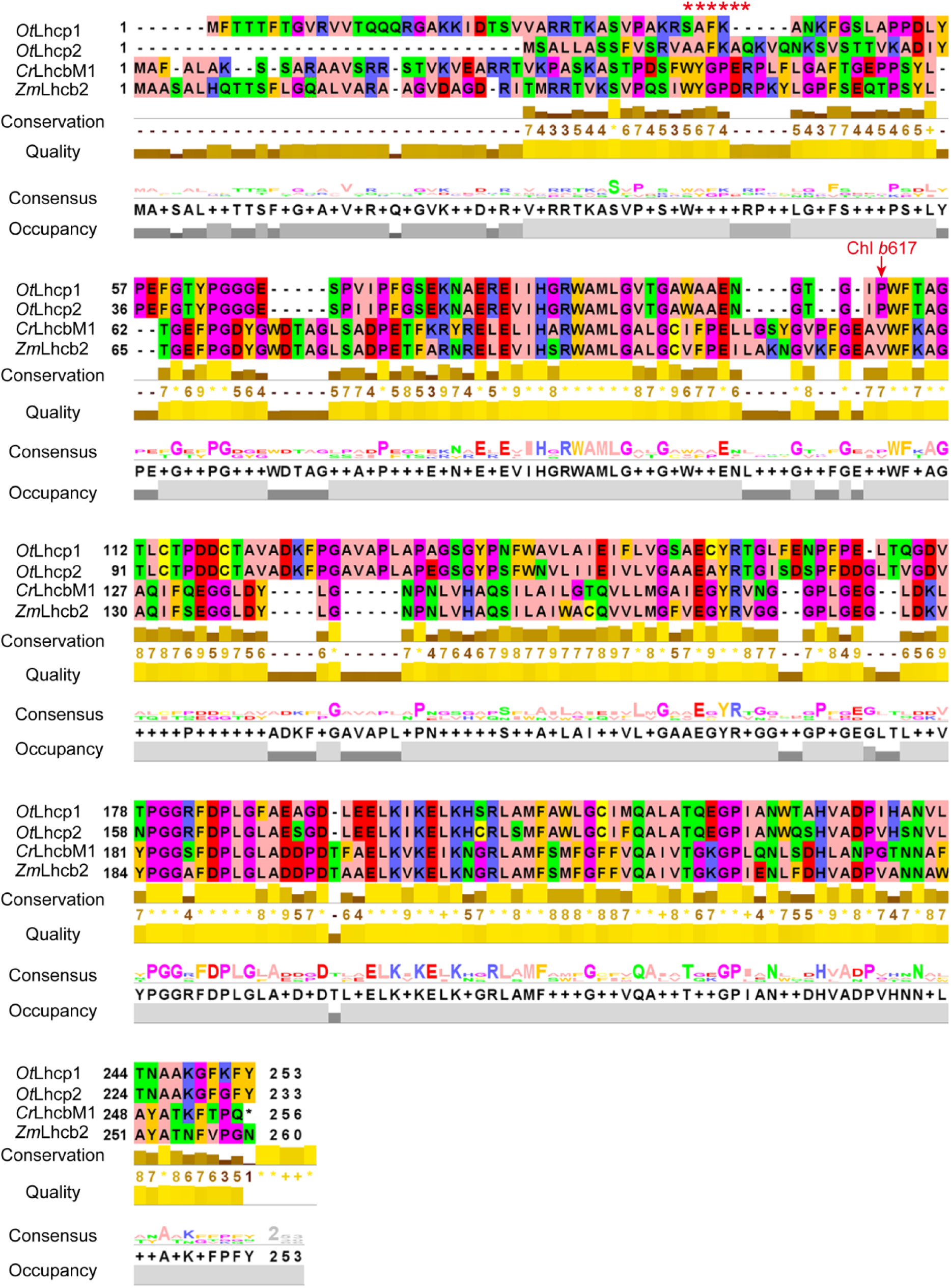
Sequence alignment of *Ot*Lhcp1 and *Ot*Lhcp2 with LhcbM1 from *C. reinhardtii* and Lhcb2 from *Z. mays.* The red arrow indicates the binding site of Chl *b*617 found in *Ot*Lhcp1 a nd Lhcp2. The red asterisk symbols show the position of WYXXXR motif found at the N-terminal regions of *Cr*LhcbM1 and *Zm*Lhcb2, but absent in *Ot*Lhcp1 and *Ot*Lhcp2.

**Figure 8—figure supplement 2.**
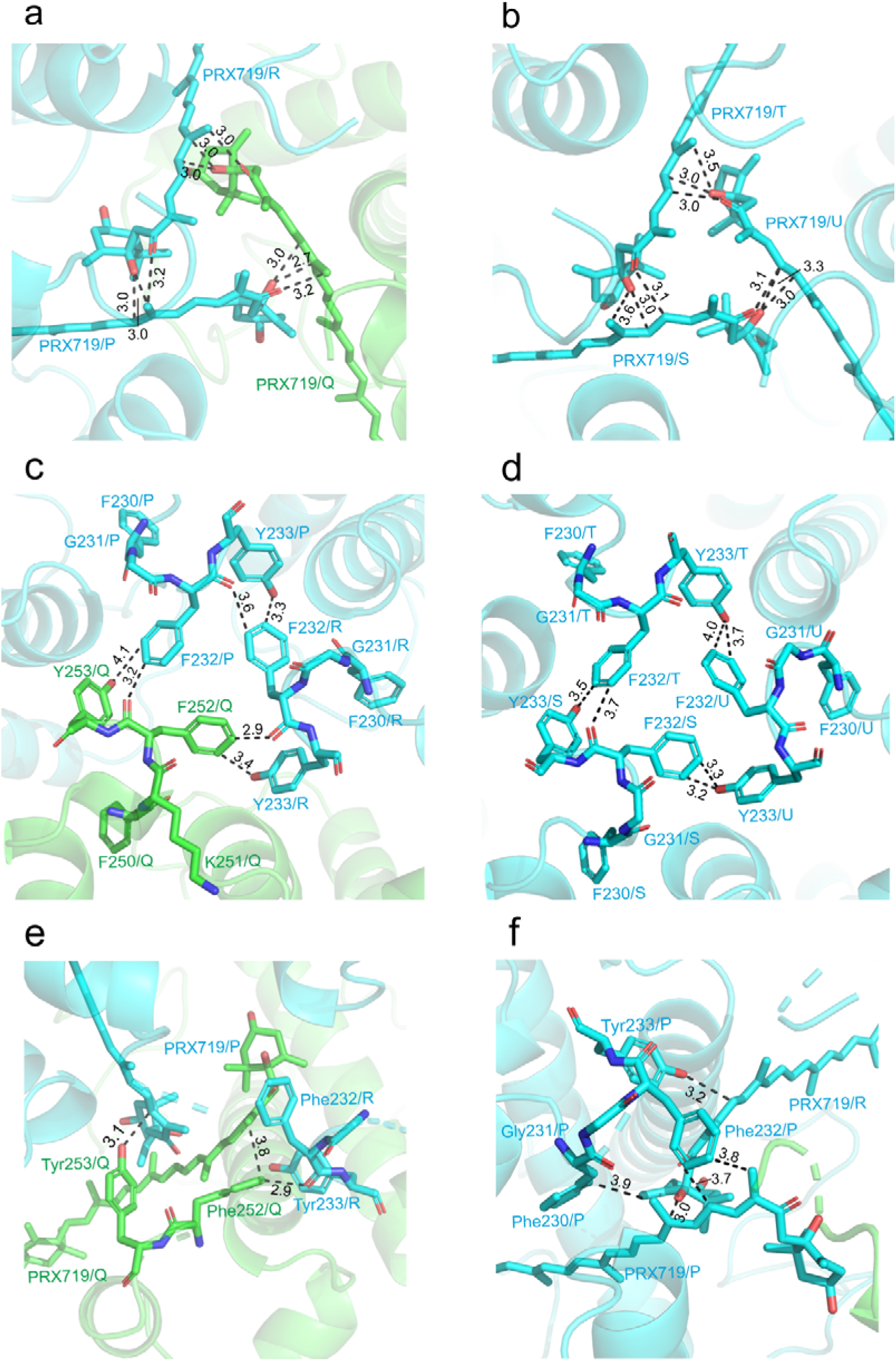
The detailed features at the trimerzation interfaces of *Ot*Trimers. **a and b**, Van der Waals and hydrophobic interactions among the three symmetry-related Prx719 molecules in the Lh cp1-(Lhcp2)_2_ trimer (a) and (Lhcp2)_3_ trimer (b); **c and d**, Interactions among the adjacent C-terminal motifs (FKFY/FGFY) of the Lhcp1-(Lhcp2)_2_ trimer (c) and (Lhcp2)_3_ (d) trimer. **e**, **f**, Extensive interactions between the C-terminal regions of *Ot*Lhcp1(e) or *Ot*Lhcp2(f) and Prx719 from the adjacent monomers. The cartoon representations of *Ot*Lhcp1 (green) and *Ot*Lhcp2 (cyan) are shown, while the molecules and amino acid residue s involved in trimerization are presented as stick models. The numbers labeled nearby the dashed lines indicate the distances (Å) between two adjacent groups.

**Figure 10—figure supplement 1.**
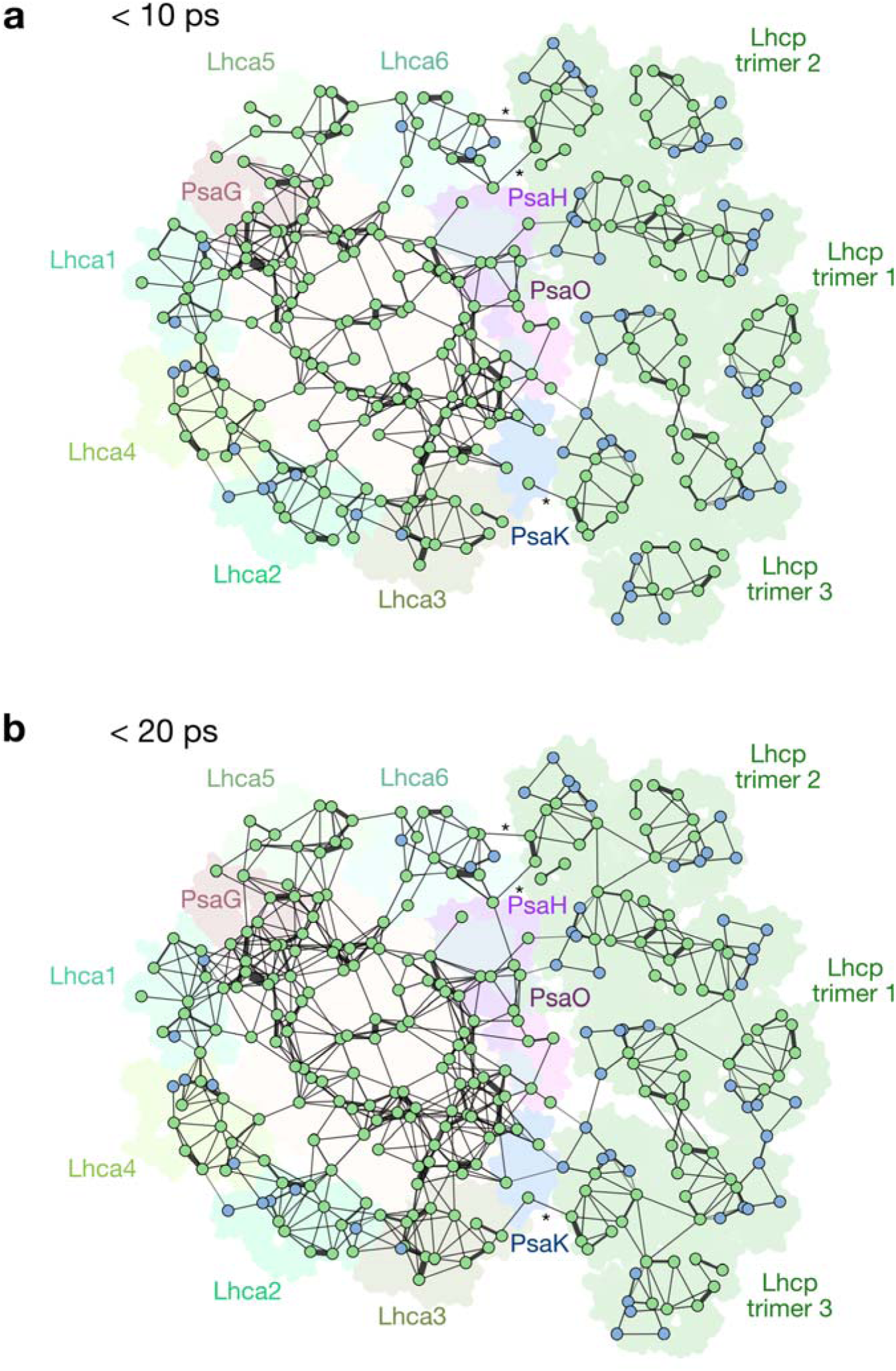
Structure-based analysis of FRET networks within the PSI-LHC- Lhcp supercomplex. a,b, Organization of Chl molecules within the PSI-LHCI-LHcp supercomplex and the FRET networks presenting efficient FRET processes with lifetimes of less than 10 ps (**a**) and 20 ps (**b**). The views are from the stromal side. The spheres and dotted circles represent each individual Chl molecules and the boundaries for the individual complexes, respectively. The lines between spheres represent the presence of FRET process between adjacent Chls. The width of lines represents the relative FRET rate in each network. The internal part of the PSI core is colored in *wheat*, while the peripheral subunits are highlighted in different colors; PsaG (*brown*), PsaH (*purple*), PsaK (*blue*), PsaO (*pink*), Lhca1-6 and Lhcp1-2 (*various greenish colors*).

**Figure 11—figure supplement 1.**
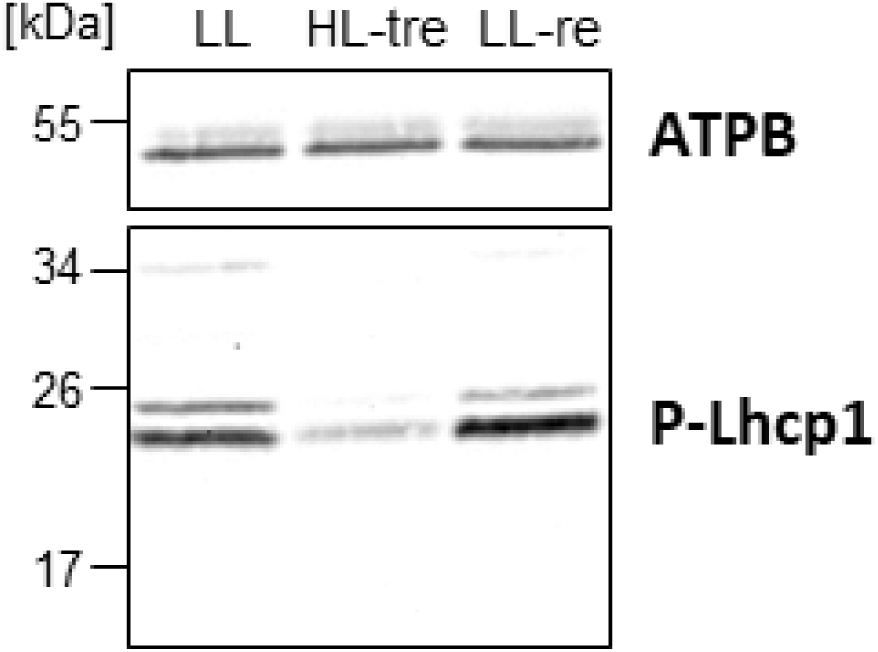
Immunoblot analysis of phospho-Lhcp1. All conditions were the same as in Fig.11b.

**Figure 11—figure supplement 2.**
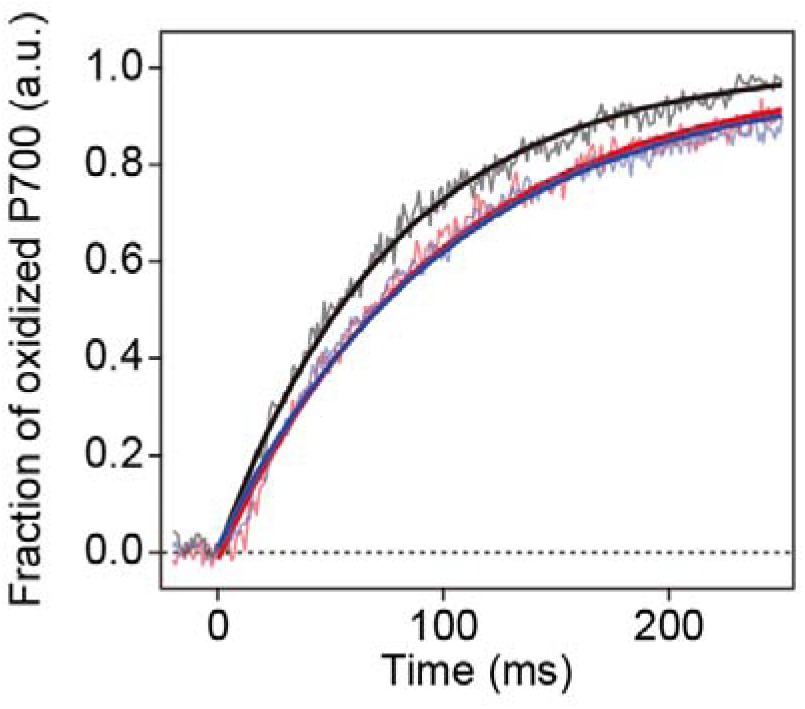
Light-induced P700 oxidation kinetics measured under 22 µmol photon m^-2^s^-1^. All conditions were the same as in Fig.11c.

**Figure 1—table supplement 1.** Pigment composition in the A3L fraction as revealed by UPLC analysis. Mean (±STD, n=3)

| Pigment | Molar ratio (Chl *a* = 100) |
| --- | --- |
| Chl *b* | 36.2 (±0.1) |
| β-carotene | 13.4 (±0.5) |
| Mdp | 3.8 (±0.1) |
| Uriolide<sup>1</sup> | 4.9 (±0.1) |
| Prx | 11.8 (±0.1) |
| Nex | 3.6 (±0.1) |
| Vio | 3.3 (±0.1) |
| Micromonal<sup>1</sup> | 2.3 (±0.1) |
| Dlt | 2.2 (±0.1) |

**Figure 1—table supplement 2.** Two-dimensional classification of the PSI particles in the A3 and A3L fractions. The RELION 2.1 package (Kimanius et al. 2016) was used for automated particle picking of particles and 2D classification into 50 classes as previously described (Watanabe et al 2019). Classes of poor quality due to aggregation, contamination, micrograph edge, or extreme proximity were discarded. Two-dimensional classification was performed on the PSI particles and manually assigned to respective small and large PSI supercomplexes.

| Fraction | A3 | A3L |
| --- | --- | --- |
| Small PSI (PSI–LHCI) | 2675 (100%) | 1030 (30%) |
| Large PSI (PSI–LHCI–Lhcp) | N/D | 2437 (70%) |
| Total PSI particles | 2675 | 3467 |

**Figure 2—table supplement 1.** Statistics of structural analysis of the *Ot*PSI-LHCI-Lhcp supercomplex.

|  | *Ot*PSI-LHCI-Lhcp |
| --- | --- |
| **Data collection and processing** |  |
| Magnification | 130,000 |
| Voltage (kV) | 300 |
| Electron exposure (e-/Å<sup>2</sup>) | 60 |
| Defocus range (μm) | from -1.8 to -2.2 |
| Pixel size (Å) | 1.04 |
| Symmetry imposed | C1 |
| Initial particle images (no.) | 5,288,217 |
| Final particle images (no.) | 80,366 |
| Map resolution (Å) | 2.94 |
| FSC threshold | 0.143 |
| Map resolution range (Å) | 2.3-4.3 |
| **Refinement** |  |
| Initial model used (PDB code) | 5ZJI, 7D0J |
| Model resolution (Å) |  |
| FSC threshold | 0.5 |
| Map sharpening B factor (Å<sup>2</sup>) | -110.961 |
| Model composition |  |
| Nonhydrogen atoms | 65071 |
| Protein residues | 5647 |
| Ligands | 449 |
| B factor (Å<sup>2</sup>) |  |
| Protein | 30.03 |
| Ligand | 39.61 |
| R.m.s. deviations |  |
| Bond lengths (Å) | 0.005 |
| Bond angles (°) | 0.973 |
| Validation |  |
| MolProbity score | 2.49 |
| Clash score | 19.9 |
| Poor rotamers (%) | 0.02 |
| Ramachandran plot |  |
| Favored (%) | 95.39 |
| Allowed (%) | 4.43 |
| Outliers (%) | 0.18 |

**Figure 2—table supplement 2.**
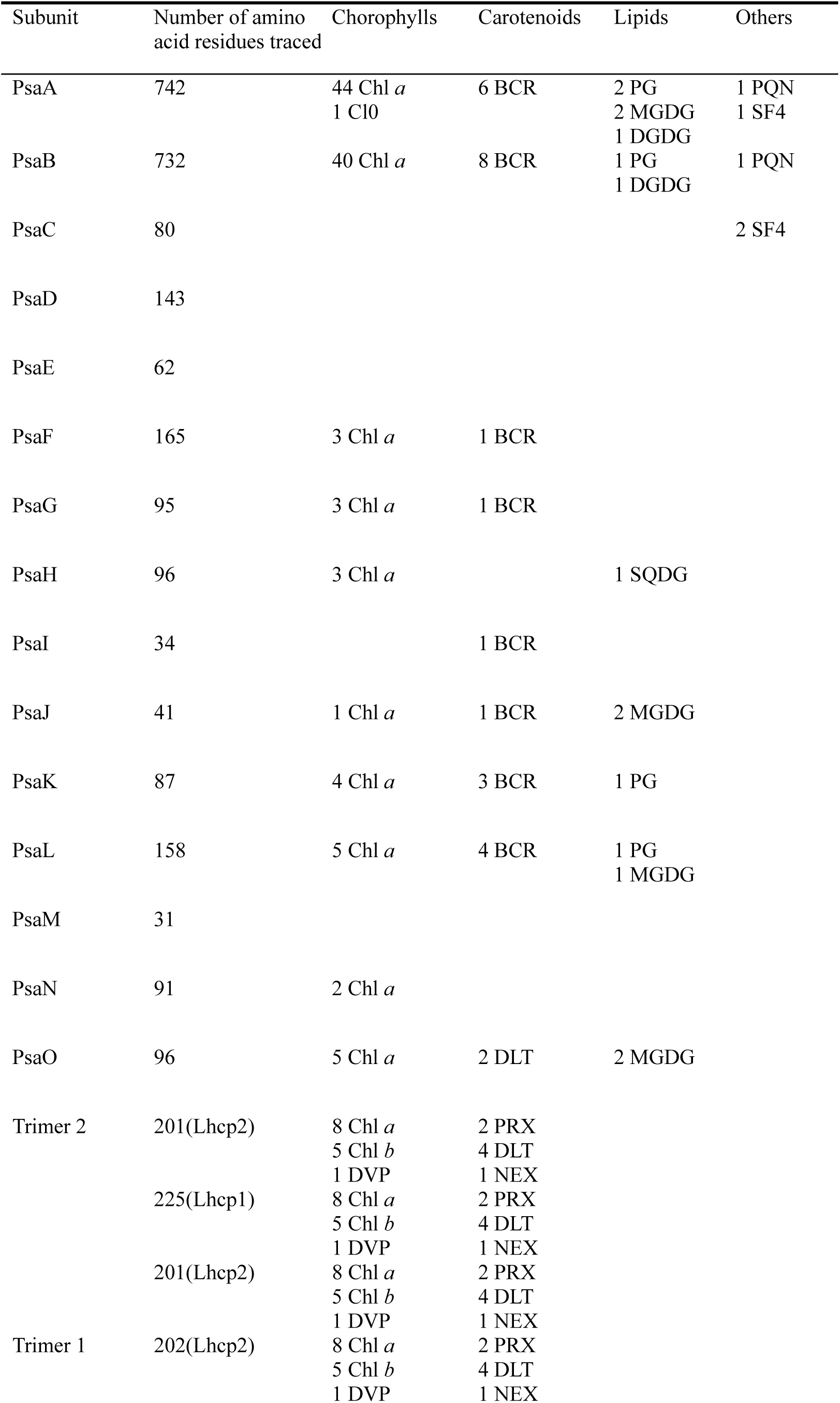
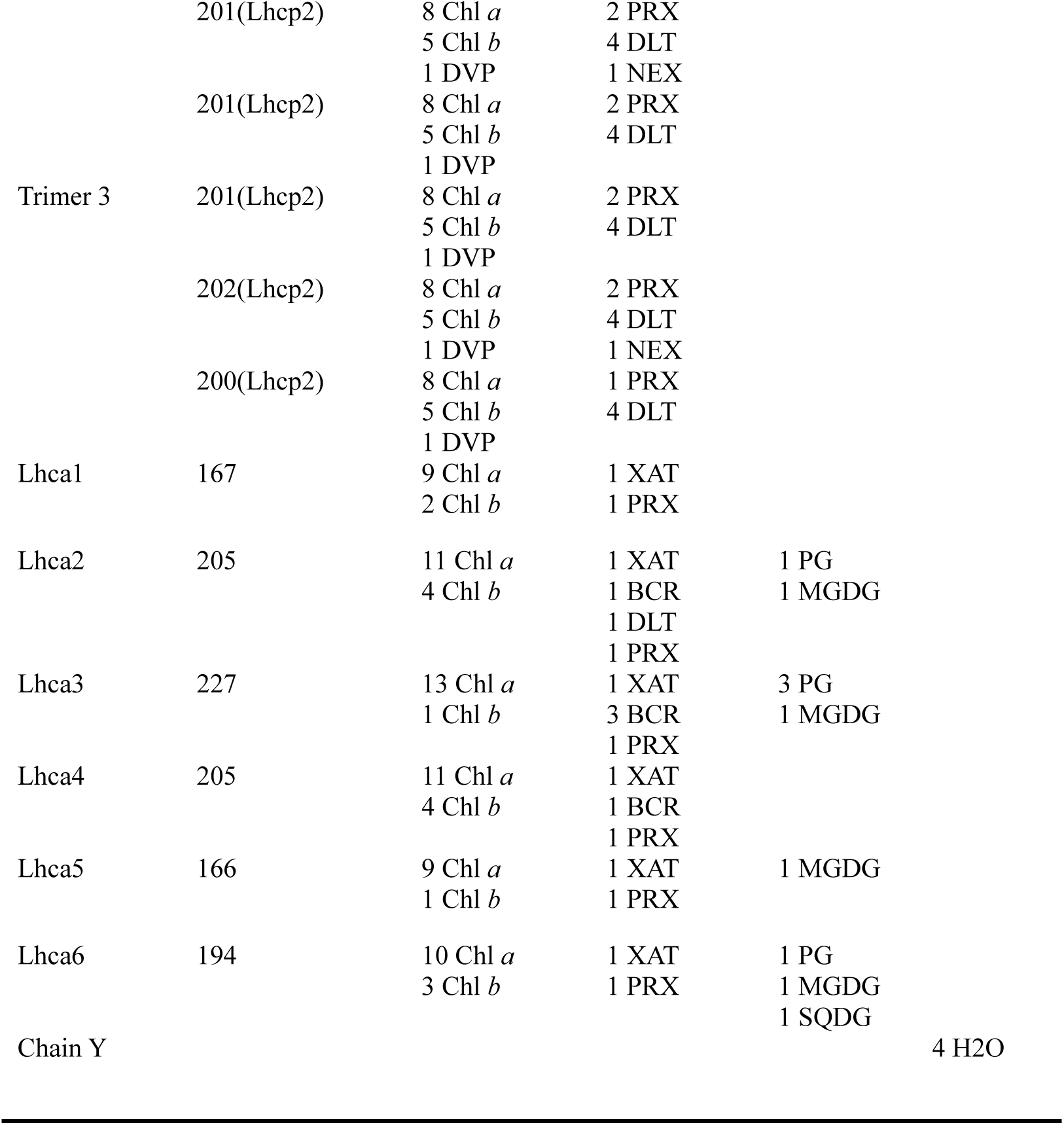
Summarization of the components in the final structural model of the *Ot*PSI-LHCI-Lhcp supercomplex.

